# IOBR: Multi-omics Immuno-Oncology Biological Research to decode tumor microenvironment and signatures

**DOI:** 10.1101/2020.12.14.422647

**Authors:** Dongqiang Zeng, Zilan Ye, Guangchuang Yu, Jiani Wu, Yi Xiong, Rui Zhou, Wenjun Qiu, Na Huang, Li Sun, Jianping Bin, Yulin Liao, Min Shi, Wangjun Liao

**Affiliations:** Department of Oncology, Nanfang Hospital, Southern Medical University, Guangzhou, Guangdong, P. R. China; Department of Bioinformatics, School of Basic Medical Sciences, Southern Medical University, Guangzhou 510515, China; Department of Neurosurgery, Xiangya Hospital, Central South University, 87 Xiangya RD, Kaifu District, Changsha 410008, China; Xiangya School of Medicine, Central South University; Department of Cardiology, Nanfang Hospital, Southern Medical University, Guangzhou, Guangdong, P. R. China

**Keywords:** tumor microenvironment, multi-omics, gene signatures, immune-tumor interaction

## Abstract

**Motivation:** Recent advance in next generation sequencing has triggered the rapid accumulation of publicly available multi-omics datasets. The application of integrated omics to exploring robust signatures for clinical translation is increasingly highlighted, attributed to the clinical success of immune checkpoint blockade in diverse malignancies. However, effective tools to comprehensively interpret multi-omics data is still warranted to provide increased granularity into intrinsic mechanism of oncogenesis and immunotherapeutic sensitivity.

**Results:** We developed a computational tool for effective Immuno-Oncology Biological Research (IOBR), providing comprehensive investigation of estimation of reported or user-built signatures, TME deconvolution and signature construction base on multi-omics data. Notably, IOBR offers batch analyses of these signatures and their correlations with clinical phenotypes, lncRNA profiling, genomic characteristics and signatures generated from single-cell RNA sequencing data in different cancer settings. Additionally, IOBR also integrates multiple existing microenvironmental deconvolution methodologies and signature construction tools for convenient comparison and selection. Collectively, IOBR is a user-friendly tool, to leverage multi-omics data to facilitate immuno-oncology exploration and unveiling of tumor-immune interactions and accelerating precision immunotherapy.

## Introduction

Clinical success of immune checkpoint blockade (ICB) has recently witnessed immunotherapy revolutionizing the treatment paradigm of advanced cancers. However, the heterogeneous immunotherapy outcomes across patients necessitates the investigation into host-tumor interactions, especially the immune cell infiltrations within tumor microenvironment (TME), to define robust predictive biomarkers for precision therapy. In this regard, increasing TME-relevant gene signatures have been reported to estimate immune contexture and predict clinical treatment response. Notably, gene expression profiling (GEP)(Cristescu, et al., 2018) and TMEscore(Zeng, et al., 2019) are influential pan-cancer predictive signatures for prognosis, ICB response and resistance by decoding TME component using transcriptomic data. Gene signatures for chemotherapy response prediction are also reported: the 70-gene(van ‘t Veer, et al., 2002) and 21-gene(Sparano and Paik, 2008) assay predict distant recurrence of estrogen receptor positive breast cancer with adjuvant chemotherapy; and aforementioned TMEscore are also promising biomarker for chemotherapy sensitivity in late-stage gastric cancer(Zeng, et al., 2019). Signatures such as PAM50, constructed by integrating transcriptomics with other omics (genomics, methylation, proteomics) to define subgroups, provide a new lens into tumor plasticity and heterogeneity of breast cancer(Cancer Genome Atlas, 2012). The emergence of these promising signatures is attributed greatly to the development of next generation sequencing (NGS) and computational deconvolution methodology. Technological breakthrough in NGS has driven enormous accumulation of publicly available multi-omics datasets, allowing easy accessibility for multi-omics data. Despite the rapid technological progress of single-cell RNA sequencing (RNA-seq), the lack of large datasets indicates that the validation of signatures still depends heavily on attainable bulk RNA-seq datasets. Additionally, based on transcriptomic data, recent development of computational algorithms and tools were utilized to dissect the tumors-microenvironment interaction. Tools for TME deconvolution is fundamentally classified according to four computational principles: machine learning, gene set enrichment analysis (GSEA), linear regression and nonlinear programming(Zhang, et al., 2020). Nonlinear programming-based principles do not necessarily rely on the information of different cell-type frequencies, whereas the other three counterparts require prior knowledge of marker genes of distinct immune cell subsets and molecular profiles(Zhang, et al., 2020). Machine learning based principles could evaluate the absolute proportion of infiltrating immune cells within TME, while gene set enrichment analysis-based principles infers the relative proportion(Zhang, et al., 2020).

Given the merits of above deconvolution methods, further comparison of the results to add accuracy and the subsequent downstream analyses are not covered by either of these tools. Competent tools to conveniently interpret transcriptomics or integrated omics data is warranted to offer new insight into tumorigenesis, immune-tumor interaction and therapeutic sensitivity diversity. Therefore, we developed a computational tool for effective Immuno-Oncology biological Research (IOBR), to comprehensively explore and visualize the multi-omics interpretation including signature score calculation and systematic estimation of its correlations with clinical phenotypes, noncoding RNA characteristics, signatures derived from single-cell RNA-seq data, and genomic landscapes in multiple cancers, as well as TME deconvolution with diverse algorithms and fast signature construction. IOBR is an effective tool and its implementation in immuno-oncology study may aid the discovery of novel tumor-immune interactions and accelerating precision immunotherapy.

## Results

To comprehensively leverage the transcriptomic data to detect immune-tumor interplay and its promising clinical translation, we introduce IOBR R package, an effective and flexible tool, freely available in the GitHub repository (https://github.com/IOBR/IOBR). IOBR consists of four functional modules, comprising estimation of signature scores, phenotype related signatures and corresponding genes, and signatures generated from single-cell RNA-seq data, along with decoding immune contexture (signature and TME deconvolution module); identification of phenotype relevant signatures, cell fraction, or signature genes, as well as pertinent batch statistical analyses (phenotype module); analysis of signature associated mutations (mutation module) and fast model construction (model construction module). The schematic workflow and functional codes are illustrated in **Figure 1** and **Figure 2**, respectively. Corresponding figures dynamically generated following inputting function-specific parameters of pertinent module. Details of each module are described below.

**Figure 1.**
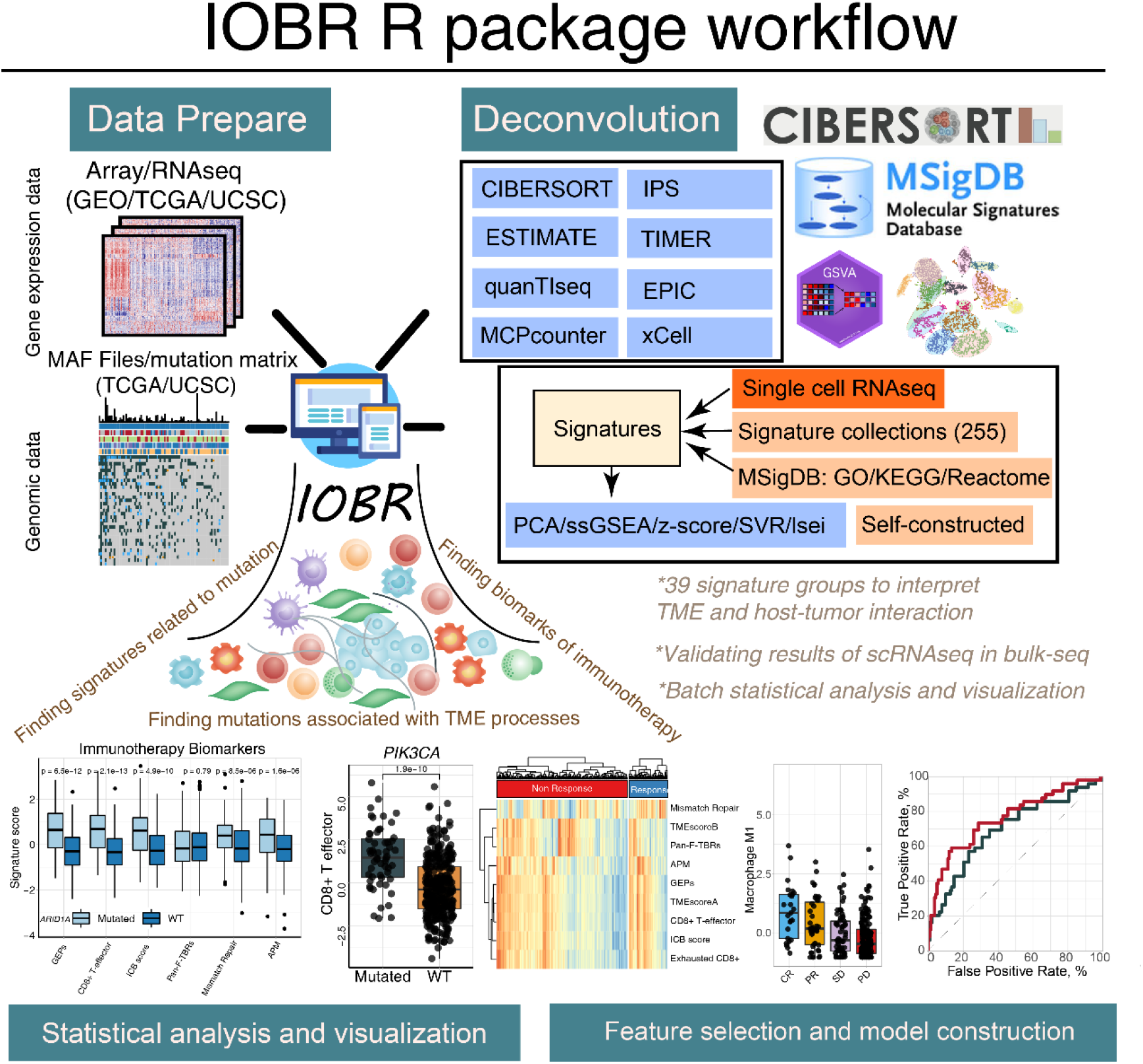
Graphical abstract outlines the workflow of IOBR package. The IOBR R package contains corresponding data preparation, multiple deconvolution algorithms to the decode signature estimation, TME contexture, batch statistical analyses and visualization, as well as feature selection and model construction.

**Figure 2.**
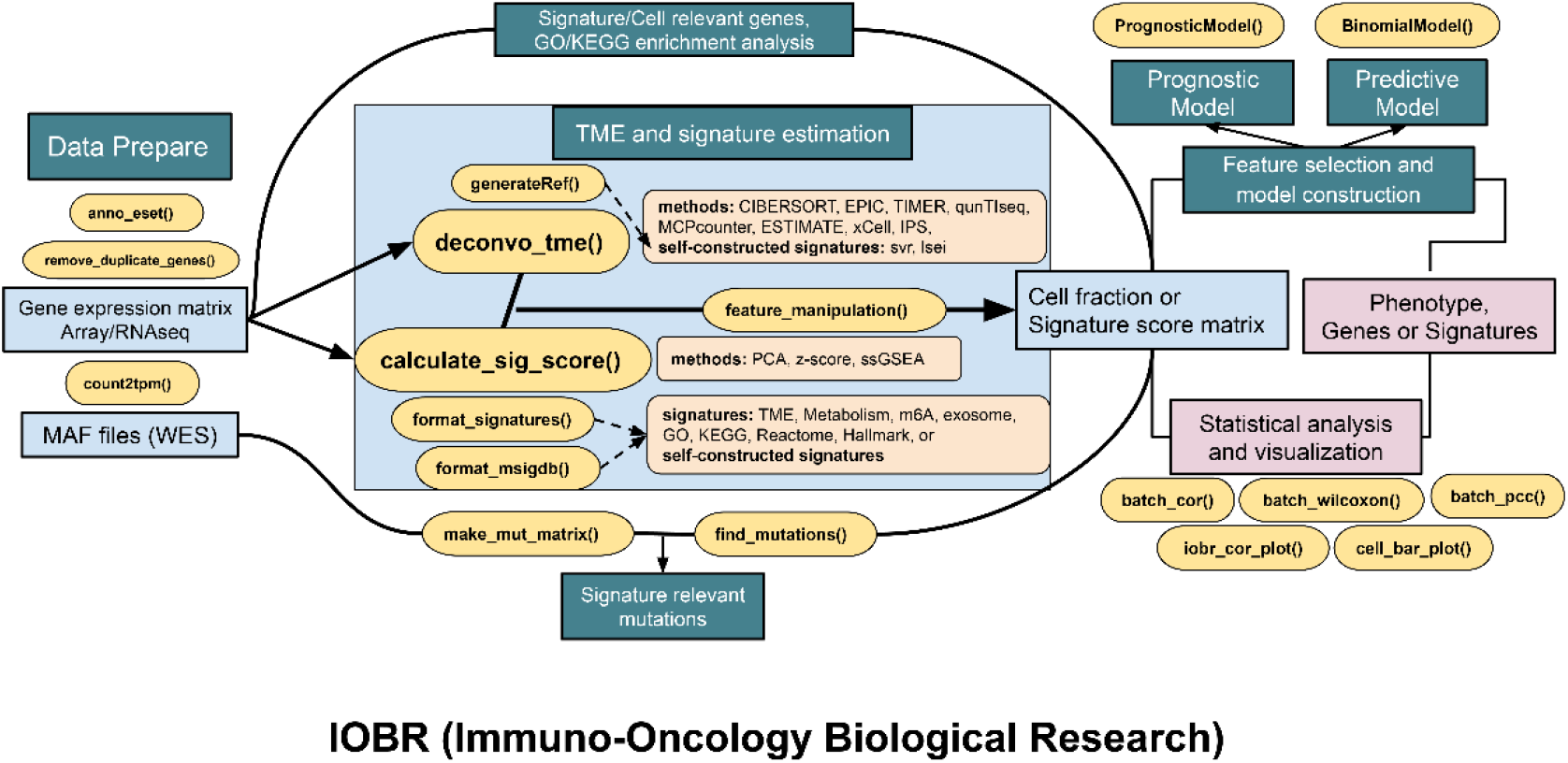
Pipeline diagram depicts functions of four analytic modules contained in IOBR. In addition to functions for data preprocessing; the function modules comprise (1) analyses of signatures pertinent to clinical phenotype, lncRNA, and targeted signatures constructed based on bulk RNA-seq or single-cell RNA-seq data and TME deconvolution; (2) identification of phenotype relevant signatures, cell fraction, or signature genes, as well as corresponding batch statistical analyses and visualization; (3) estimation of specific mutation landscape associated with interested signature (4) and model construction following feature selection.

### Signature and TME estimation module

#### Signature estimation

To elucidate an increasingly granular view of the tumor microenvironment cellular composition and functional status with the goal of cancer-therapy refinement, we construct estimation function for user-generated signatures or 255 reported signatures enrolled in IOBR. The extensive signature collection is classified into 3 categories: TME-associated, tumor-metabolism, and tumor-intrinsic signatures. Additionally, IOBR supports estimation of the signature gene sets derived from GO, KEGG, HALLMARK and REACTOME database. To note, IOBR permits user to generate a signature list based on their own biological discovery or expletory requirement, for convenient estimation and follow-up systematic exploration.

Three methodologies were included in the process of signature score evaluation, comprising Single-sample Gene Set Enrichment Analysis (ssGSEA), Principal component analysis (PCA), and Z-score. ssGSEA, is a wildly-adopted tool to calculate separate enrichment scores for each pairing of a sample and gene set(Barbie, et al., 2009). Each ssGSEA enrichment score represents the degree to which the genes in a particular gene set are coordinately up- or down-regulated within a sample. Notably, PCA computes the principal components to perform a change of basis on the exploratory data analysis for predictive model construction. Current signatures constructed using PCA methodology includes the Pan-F-TBRs(Mariathasan, et al., 2018) and the TMEscore (Zeng, et al., 2019), two promising biomarkers to predict clinical outcome and therapy sensitivity of malignancies. Z-score is a numerical measurement to describe a score’s relationship to the mean of a group of values. Z-score is measured in terms of standard deviations from the mean. These three methods are selectable in IOBR by inputting targeted methods or integration of them, and corresponding visualizations are supported.

#### Signatures derived from single-cell RNA-seq Data

Technological and computational innovations of single-cell analysis make it a popular alternative to determine cell markers and gene signatures for phenotypes. However, the significantly expensive cost and high requirement for starting tumor material limits its widespread utility. Therefore, the application of large and attainable bulk RNA-seq datasets is still a major source to validate the clinical significance of the signatures generated from single-cell analysis. Therefore, IOBR provides multiple methodologies to extract cell signature genes from the single-cell RNA-seq data (TPM or counts data are available inputs). Remarkedly, the linear support vector regression (SVR) algorithm of CIBERSORT or LSEI(Gong and Szustakowski, 2013) algorithm is implemented in IOBR for convenient bulk RNA-seq data analysis to verify the clinical value of the targeted cells identified by single-cell RNA-seq data.

#### TME deconvolution

Clinical investigations have highlighted cell infiltrations in TME as pivotal contributors to the complex anti-tumor immunity in malignancies. TME-cell deconvolution is the major technological hurdle and the deconvolution algorithms vary in merits and pitfalls. IOBR integrates eight open-source deconvolution methodologies, including CIBERSORT(Newman, et al., 2015), ESTIMATE(Yoshihara, et al., 2013), quanTIseq(Finotello, et al., 2019), TIMER(Li, et al., 2017), IPS(Charoentong, et al., 2017), MCPCounter(Becht, et al., 2016), xCell(Aran, et al., 2017) and EPIC(Racle, et al., 2017).

CIBERSORT is the most well-recognized method to detect 22 immune cells in TME, allowing large-scale analysis of RNA mixtures for cellular biomarkers and therapeutic targets with promising accuracy(Newman, et al., 2015). Notably, IOBR, adopting the linear vector regression principle of CIBERSORT, allows users to construct self-defined signature. The availability of its input file was extended to cell-subset derived from single-cell sequencing results. ESTEMATE dissects non-malignant contexture including stromal and immune signatures to determine tumor purity(Yoshihara, et al., 2013). The method of quanTIseq enumerates ten immune cell subsets from bulk RNAseq data(Finotello, et al., 2019). TIMER quantifies the abundance of six tumor-infiltrating immune compartments and firstly provides 6 major analytic modules to analyzed the immune infiltration with other cancer molecular profiles(Li, et al., 2017). IPS estimates 28 TIL subpopulations including effector and memory T cells and immunosuppressive cells(Charoentong, et al., 2017). MCP-counter conducts robust quantification of the absolute abundance of eight immune and two stromal cell populations in heterogeneous tissues from transcriptomic data(Becht, et al., 2016). xCell provides comprehensive view of 64 immune cells from RNA-seq data and other cell subsets in bulk tumor tissue(Aran, et al., 2017). EPIC refers the proportion of immune and cancer cells from the expression of genes and compare it with the gene expression profiles from specific cells to predict the cell subpopulation landscape(Racle, et al., 2017). In a nutshell, IOBR allows convenient integration and visualization of above deconvolution results or flexible selection of particular methodology of interest.

### Phenotype module

To implement above TME deconvolution and signatures calculation to explore potential clinical translation, we collect and systematically categorize the signatures into 39 groups. The categories involve TME cell populations (classified either by deconvolution methods or cell types), signatures of immunophenotype, tumor metabolism, hypoxia, and EMT *et al*. Furthermore, IOBR supports constructing a novel signature group derived from their own immuno-oncological findings to lay foundation for subsequent minding latent biological mechanism and potential clinical translation. Notably, the “iobr_cor_plot” function is included into IOBR, to dynamically generate statistical results and to efficaciously depict the correlation between signatures and targeted phenotype, such as therapeutic responses, carcinogenic infection status. Additionally, IOBR is capable of quick visualizing the relationships between signature genes and the targeted variable (binary or continuous) with identical methods. Likewise, IOBR is also feasible to identify signatures significantly correlated with the signature of interest.

Moreover, the “iobr_cor_plot” function is also effectively available to define the signatures correlated with long non-coding RNA (lncRNA) profiling, by extracting targeted gene from the lncRNA expression matrix as a phenotype. The subsequent batch correlation analysis procedure is similar to priorly described. Additionally, in that pertinent signatures and signature genes could be multiple, IOBR enrolled a subset of functions for batch statistical analysis and visualization. It comprises the batch survival analysis for either continuous signature scores or categorized phenotype subgroups, and aforementioned batch correlation analysis using statistical tests including Wilcoxon test and Partial correlation coefficient (PCC) correspondingly.

Collectively, phenotype module of IOBR R package permits systematic identification of phenotype relevant signatures, cell fraction, or signature genes, as well as corresponding batch statistical analyses and visualization.

### Mutation module

In addition to systematical signature-phenotype investigation, IOBR expands the transcriptomic exploration to the interplay with genome profiles. Genome data with MAF-format(Mayakonda, et al., 2018) downloaded from University of California, Santa Cruz (UCSC) website, or user-construct mutation matrix is acceptable as an input to dig out mutations related to specific signatures. Furthermore, IOBR supports convenient transforming the MAF data into a mutation matrix with distinct variation types comprising insertion–deletion mutations (indel), single-nucleotide polymorphism (SNP), frameshift, or an integrate of them all for flexible selection. Wilcoxon ranksum test is employed in this module for batch analysis of mutations significantly associated with targeted signatures. IOBR also supports batch visualization of the mutation statutes (mutation or non-mutation) of interest.

### Model construction module

For effective application of the signatures in clinical interpretation, IOBR provides functions for feature selection, robust biomarker identification, and model construction based on priorly identified phenotype associated signatures. To our knowledge, the therapeutic response and overall survival is the focused endpoints in oncology, and leveraging the corresponding signatures to construct models might hold promise in precise and cost-effective prediction of tumor prognosis and treatment sensitivity. Moreover, rational utility in other bioscience settings may also shed new light on uncovering novel discoveries of interest.

### Application

The detailed the implementation of IOBR was illustrated in the Supplementary Materials by a complete analysis pipeline. To note, in a recent published literature with multi-omics data from IMvigor210 cohort, we generated immunotherapy associated risk score, determined TME infiltration pattern and further located in macrophage as a robust predictive biomarker, subsequently unveiled the predominant genomic alterations and significant metabolic characteristics (Zeng, et al., 2020). Charts derived from IOBR reach quality requirements of publication and can be flexibly modified locally.

## Discussion

The complexity and increasing accumulation of multi-omics datasets pose new opportunity for integrative analysis of immuno-oncology, and also challenges to simplify the interpretation without sacrificing the high accuracy. Our study developed a comprehensive computational tool IOBR to dissect host-tumor interaction and signatures for therapeutic sensitivity. Four major analytic modules were provided, allowing effective and systematical analysis of tumor immunologic, clinical, genomics, and single-cell RNA-seq data.

With the era of immunotherapy and Big data coming, identifying novel biomarkers and calculating signatures to finetune therapy strategies have come to the spotlight of immune-oncology. In addition to systematic estimation of published signature score and signature constructed by users, IOBR is competent to operate and interpretate the lncRNA profiling, gene alteration landscapes, single-cell RNA-seq results. Notably, the validation of signatures generated by single-cell analysis is also involved, which relies intensely on large bulk RNA-seq datasets. Additionally, the model construction module potentiates the innovative clinical translation of signatures genes into prediction of tumor prognosis, therapy response and resistance. Moreover, the tumor microenvironment is an essential constituent of tumor immunity, and the correlation between TME heterogenicity and clinical phenotype is pivotal for preclinical oncology research. IOBR R package offers multiple available deconvolution methodologies which removed the roadblock for decoding TME contexture. TIMER is a published web tool integrating six algorithms for inferring immune cell composition from bulk tumor transcriptome profiles(Li, et al., 2020). However, despite the convenience of intuitive outputs provided by TIMER2.0, the upload of large dataset is still a challenge for website tool, which could be tackled by R package tools to better analyze data with larger volume of samples and to convenient acquisition of large data results.

With the multi-omics data accumulation, we anticipate IOBR to attract broad application in immuno-oncology and facilitate accelerate the discovery of latent immune evasion mechanisms and novel therapeutic targets. IOBR represents a contribution to the computational toolbox for unveiling immune-tumor interactions from multi-omics data, and implementing it in preclinical researches of tumor heterogeneity and plasticity may be instrumental to provide impetus for precision immunotherapy.

## Supporting information

IOBR VIGNETTE

## Availability

https://github.com/IOBR/IOBR

## Authors’ Contributions

*Study concept and design:* Dongqiang Zeng;

*Acquisition of data:* Zilan Ye, Jiani Wu, Wenjun Qiu, Na Huang, Li Sun;

*Analysis and interpretation of data:* Dongqiang Zeng, Rui Zhou;

*Package development:* Dongqiang Zeng, Zilan Ye;

*Drafting of the tutorial:* Dongqiang Zeng, Zilan Ye;

*Drafting of the manuscript:* Zilan Ye, Dongqiang Zeng;

*Critical revision of the manuscript for important intellectual content:* Guangchuang Yu, Min Shi, Wangjun Liao;

*Obtaining funding:* Min Shi, Wangjun Liao;

*Administrative, technical, or material support:* Yi Xiong, Jianping Bin, Yulin Liao;

*Supervision:* Guangchuang Yu, Wangjun Liao;

*Other:* None

## Acknowledgments

We extend our sincere thanks to *Dr. Rongfang Shen* from Chinese Academy of Medical Sciences and Peking Union Medical College for the assistance in functions of model construction and TME deconvolution. Function ‘deconvo_tme’ is inspired from the article “Comprehensive evaluation of transcriptome-based cell-type quantification methods for immuno-oncology”. Thanks to authors for making source code available.

