## Supplementary material for "IOBR: Multi-omics Immuno-Oncology Biological Research to decode tumor microenvironment and signatures": IOBR VIGNETTE: IOBR-VIGNETTE-20201215.html

IOBR Immuno-Oncology Biological Research


### IOBR Immuno-Oncology Biological Research

###### **Dongqiang Zeng**

**Department of Oncology,   
Nanfang Hosipital,   
Southern Medical University, Guangzhou, China**  
****

#### **2020-12-15**

### CONTENTS

- 1. Introduction
- 2. Installation
- 3. Data Scenario and Preparation
  - 3.1 Gene-expression Data Derived from RNAseq
  - 3.2 Gene-expression Data Derived from Array
- 4. IOBR Pipeline Introduction
- 5. Functional Modules
  - 5.1 Signature And TME Deconvolution Module
    - 5.1.1 Signature Estimation
      - 1) Signature collection
      - 2) Methods for signature calculation
    - 5.1.2 Signatures Derived From Single Cell RNAseq
      - 1) Methods for signature estimation
      - 2) Signature-phenotype correlation
    - 5.1.3 Available Methods to Decode TME Contexture
      - 1) Method 1: CIBERSORT
      - 2) Method 2: EPIC
      - 3) Method 3: MCPcounter
      - 4) Method 4: xCELL
      - 5) Method 5: ESTIMATE
      - 6) Method 6: TIMER
      - 7) Method 7: quanTIseq
      - 8) Method 8: IPS
      - 9) Combination of above deconvolution results
  - 5.2 Phenotype Module
    - 5.2.1 Explore Phenotype Relevant Signatures
      - 1) Load data
      - 2) Signature groups
      - 3) Identify phenotype relevant signatures
      - 4) Visualize phenotype relevant signature genes
      - 5) Estimate lncRNA associated signatures
    - 5.2.2 Identify Signatures Relevant to Targeted Signature
      - 1) Construct phenotype group
      - 2) Analyze Pan-F-TBRs correlated signatures
      - 3) Evaluate Pan-F-TBRs related TME cell infiltration
    - 5.2.5 Batch Analyses and Visualization
      - 1) Batch survival analyses
      - 2) Subgroup survival analyses
      - 3) Batch statistical analyses
      - 4) Batch analyses of Pearson’s correlation coefficient
  - 5.3 Mutation Module
    - 5.3.1 Load Mutation MAF data
    - 5.3.2 Analyze Signature Associated Mutations
  - 5.4 Model Construction Module
- 6. Summary
- 7. Session Information
- 8. Citing IOBR
- REFERENCES

### 1. Introduction

Recent advance in next generation sequencing has triggered the rapid accumulation of publicly available multi-omics data^1^. The application of integrated omics to exploring robust signatures for clinical translation is increasingly highlighted in immuno-oncology,but raises computational and biological challenges^2^. This vignette aims to demonstrate how to utilize the package named IOBR to perform multi-omics immuno-oncology biological research to decode tumor microenvironment and signatures for clinical translation. This R package integrates 8 published methodologies for decoding tumor microenvironment (TME) contexture: `CIBERSORT`, `TIMER`, `xCell`, `MCPcounter`, `ESITMATE`, `EPIC`, `IPS`, `quanTIseq`. Moreover, 255 published signature gene sets were collected by IOBR, involving tumor microenvironment, tumor metabolism, m6A, exosomes, microsatellite instability, and tertiary lymphoid structure. Run the function `signature_collection_citation` to attain the source papers and the function `signature_collection` returns the detail signature genes of all given signatures. Subsequently, IOBR adopts three computational methods to calculate the signature score, comprising `PCA`,`z-score`, and `ssGSEA`. To note, IOBR collected and employed multiple approaches for variable transition, visualization, batch survival analysis, feature selection, and statistical analysis. Batch analysis and visualization of corresponding results is supported. The details of how IOBR works is described below.

##### Graphical abstract outlines the workflow of IOBR package.

### 2. Installation

It is essential that you have R 3.6.3 or above already installed on your computer or server. IOBR is a pipeline that utilizes many other R packages that are currently available from CRAN, Bioconductor and GitHub.

Before installing IOBR, please properly install all dependencies including `tibble`, `survival`, `survminer`, `limma`, `limSolve`, `GSVA`, `e1071`, `preprocessCore`, `ggplot2`, `ggpubr` and so on. It is an easy way to install them by executing the following command in your R session:

```
# options("repos"= c(CRAN="https://mirrors.tuna.tsinghua.edu.cn/CRAN/"))
# options(BioC_mirror="http://mirrors.tuna.tsinghua.edu.cn/bioconductor/")
if (!requireNamespace("BiocManager", quietly = TRUE)) install("BiocManager")
depens<-c('tibble', 'survival', 'survminer', 'sva', 'limma', "DESeq2","devtools",
          'limSolve', 'GSVA', 'e1071', 'preprocessCore', 'ggplot2', "biomaRt",
          'ggpubr', "devtools", "tidyHeatmap", "caret", "glmnet", "ppcor","timeROC","pracma")
for(i in 1:length(depens)){
  depen<-depens[i]
  if (!requireNamespace(depen, quietly = TRUE)) BiocManager::install(depen)
}

if (!requireNamespace("EPIC", quietly = TRUE))
  devtools::install_github("GfellerLab/EPIC", build_vignettes=TRUE)
if (!requireNamespace("MCPcounter", quietly = TRUE))
  devtools::install_github("ebecht/MCPcounter",ref="master", subdir="Source")
if (!requireNamespace("estimate", quietly = TRUE)){
  rforge <- "http://r-forge.r-project.org"
  install.packages("estimate", repos=rforge, dependencies=TRUE)
}
```

Then, you can start to install IOBR from github by typing the following code into your R session:

```
if (!requireNamespace("remotes", quietly = TRUE)) install("remotes")
if (!requireNamespace("IOBR", quietly = TRUE))
  remotes::install_github("IOBR/IOBR",ref="master")
```

Load the IOBR package in your R session after the installation is complete:

```
library(IOBR)
library(EPIC)
library(estimate) 
library(MCPcounter)
library(tidyverse)
library(tidyHeatmap)
library(maftools)
library(ggpubr)
library(ggplot2)
library(survival)
```

### 3. Data Scenario and Preparation

#### 3.1 Gene-expression Data Derived from RNAseq

##### Preparation of Gene-expression data derived from RNAseq.

For transcriptomic data of TCGA data sets, we strongly recommend user to use UCSCXenaTools R package. Here, we download counts data of TCGA-STAD from UCSC using UCSCXenaTools R package. If you use it in published research, please cite: *Wang et al.*, (2019). The UCSCXenaTools R package: a toolkit for accessing genomics data from UCSC Xena platform, from cancer multi-omics to single-cell RNA-seq. Journal of Open Source Software, 4(40), 1627, https://doi.org/10.21105/joss.01627. If you are not online or can not access to UCSC, we have prepared expression data of TCGA-STAD in IOBR package.`data(eset_stad)` could be applied to loading gene expression data.

```
if (!requireNamespace("UCSCXenaTools", quietly = TRUE))
    BiocManager::install("UCSCXenaTools")
library(UCSCXenaTools)
#> =========================================================================================
#> UCSCXenaTools version 1.3.4
#> Project URL: https://github.com/ropensci/UCSCXenaTools
#> Usages: https://cran.r-project.org/web/packages/UCSCXenaTools/vignettes/USCSXenaTools.html
#> 
#> If you use it in published research, please cite:
#> Wang et al., (2019). The UCSCXenaTools R package: a toolkit for accessing genomics data
#>   from UCSC Xena platform, from cancer multi-omics to single-cell RNA-seq.
#>   Journal of Open Source Software, 4(40), 1627, https://doi.org/10.21105/joss.01627
#> =========================================================================================
#>                               --Enjoy it--
# NOTE: This process may take a few minutes which depends on the internet connection speed. Please wait for its completion.
eset_stad<-XenaGenerate(subset = XenaCohorts =="GDC TCGA Stomach Cancer (STAD)") %>% 
  XenaFilter(filterDatasets    = "TCGA-STAD.htseq_counts.tsv") %>% 
  XenaQuery() %>%
  XenaDownload() %>% 
  XenaPrepare()
#> This will check url status, please be patient.
#> All downloaded files will under directory /tmp/Rtmpmi7SbB.
#> The 'trans_slash' option is FALSE, keep same directory structure as Xena.
#> Creating directories for datasets...
#> Downloading TCGA-STAD.htseq_counts.tsv.gz
eset_stad[1:5,1:5]
#> # A tibble: 5 x 5
#>   Ensembl_ID `TCGA-D7-5577-0… `TCGA-D7-6818-0… `TCGA-BR-7958-0… `TCGA-D7-8572-0…
#>   <chr>                 <dbl>            <dbl>            <dbl>            <dbl>
#> 1 ENSG00000…            11.3             10.6             10.2             10.8 
#> 2 ENSG00000…             1                0                0                2   
#> 3 ENSG00000…            11.2              9.96            10.2             11.6 
#> 4 ENSG00000…             9.13             8.77             8.73            10.1 
#> 5 ENSG00000…             8.42             8.89             8.46             9.87
```

##### Transform gene expression matrix into TPM format, and conduct subsequent annotation.

```
# Remove the version numbers in Ensembl ID.
eset_stad$Ensembl_ID<-substring(eset_stad$Ensembl_ID, 1, 15)
eset_stad<-column_to_rownames(eset_stad, var = "Ensembl_ID")
# Revert back to original format because the data from UCSC was log2(x+1)transformed.
eset_stad<-(2^eset_stad)+1

# In this process, *biomaRt* R package is utilized to acquire the gene length of each Ensembl ID and calculate the TPM of each sample. If identical gene symbols exists, these genes would be ordered by the mean expression levels. The gene symbol with highest mean expression level is selected and remove others.
# NOTE: This process may take a few minutes which depends on the internet connection speed. Please wait for its completion.
eset_stad<-count2tpm(countMat = eset_stad, idType = "Ensembl", org="hsa")
eset_stad[1:5,1:5]
#>         TCGA-D7-5577-01A TCGA-D7-6818-01A TCGA-BR-7958-01A TCGA-D7-8572-01A
#> MT-CO1          41168.66        104364.29         55792.50         71064.82
#> MT-CO3          36231.83         45850.89         36682.78         33767.40
#> MT-ND4          25999.46         44800.15         31554.70         39443.03
#> MT-CO2          33418.75         61079.13         36951.59         31441.07
#> MT-RNR2         34755.20         65345.57         26787.41         23654.63
#>         TCGA-VQ-A91Z-01A
#> MT-CO1          51781.74
#> MT-CO3          57959.58
#> MT-ND4          55387.51
#> MT-CO2          43161.95
#> MT-RNR2         49405.00
```

#### 3.2 Gene-expression Data Derived from Array

##### Obtain dataset from GEO Gastric cancer:GSE100935 using GEOquery R package.

```
if (!requireNamespace("GEOquery", quietly = TRUE))
    BiocManager::install("GEOquery")
library("GEOquery")
# NOTE: This process may take a few minutes which depends on the internet connection speed. Please wait for its completion.
eset_geo<-getGEO(GEO     = "GSE100935",
                 getGPL  = F,
                 destdir = "./")

eset    <-eset_geo[[1]]
eset    <-exprs(eset)
eset[1:5,1:5]
#>           GSM2696792 GSM2696793 GSM2696794 GSM2696795 GSM2696796
#> 1007_s_at  10.261650  10.740100  10.694980  10.692940  10.565560
#> 1053_at     8.365080   8.463314   8.535923   8.091345   7.280246
#> 117_at      7.212391   5.832635   5.616512   8.090054   4.959763
#> 121_at      7.377803   7.549826   8.186557   7.459644   7.782259
#> 1255_g_at   6.699024   2.621356   2.422049   2.423371   2.603957
```

##### Annotate genes in expression matrix and remove duplicate genes.

```
# Load the annotation file `anno_hug133plus2` in IOBR.
head(anno_hug133plus2)
#> # A tibble: 6 x 2
#>   probe_id  symbol 
#>   <fct>     <fct>  
#> 1 1007_s_at MIR4640
#> 2 1053_at   RFC2   
#> 3 117_at    HSPA6  
#> 4 121_at    PAX8   
#> 5 1255_g_at GUCA1A 
#> 6 1294_at   MIR5193
# Conduct gene annotation using `anno_hug133plus2` file; If identical gene symbols exists, these genes would be ordered by the mean expression levels. The gene symbol with highest mean expression level is selected and remove others. 

eset<-anno_eset(eset       = eset,
                annotation = anno_hug133plus2,
                symbol     = "symbol",
                probe      = "probe_id",
                method     = "mean")
eset[1:5, 1:5]
#>              GSM2696792 GSM2696793 GSM2696794 GSM2696795 GSM2696796
#> SH3KBP1        14.30308   14.56398   14.37668   14.47983   14.56702
#> RPL41          14.36095   14.32783   14.33181   14.34614   14.35773
#> LOC101928826   14.18638   14.38247   14.34530   14.26433   14.23477
#> EEF1A1         14.13245   14.15141   14.08980   14.05879   14.09277
#> B2M            14.17932   14.25636   14.01158   14.12871   14.05124
# Another annotation file `anno_illumina` is also provided by IOBR for convenient annotation of Illumina gene expression arrays  
head(anno_illumina)
#> # A tibble: 6 x 2
#>   probe_id     symbol   
#>   <fct>        <fct>    
#> 1 ILMN_1725881 LOC23117 
#> 2 ILMN_1910180 HS.575038
#> 3 ILMN_1804174 FCGR2B   
#> 4 ILMN_1796063 TRIM44   
#> 5 ILMN_1811966 LOC653895
#> 6 ILMN_1668162 DGAT2L3
```

### 4. IOBR Pipeline Introduction

##### Pipeline diagram dipict major functions contained in IOBR which are categorized into four functional modules.

!

##### Outline of all functions for data preparation and analyses of each module.

- **Data Preparation: data annotation and transformation**

  - `count2tpm()`: transform count data of RNA sequencing into TPM data.
  - `anno_eset()`: annotate the normalized genes expression matrix, including RNAseq and array (Affymetrix or Illumina).
  - `remove_duplicate_genes()`: remove the genes annotated with the duplicated symbol after normalization and retain only the symbol with highest expression level.
- **TME Deconvolution Module: integrate multiple algorithms to decode immune contexture**

  - `deconvo_tme()`: decode the TME infiltration with different deconvolution methodologies, based on bulk RNAseq, microarray or single cell RNAseq data.
  - `generateRef()`: generate a novel gene reference matrix for a specific feature such as infiltrating cell, through the SVR and lsei algorithm.
- **Signature Module: calculate signature scores, estimate phenotype related signatures and corresponding genes, and evaluate signatures generated from single-cell RNA sequencing data**

  - `calculate_sig_score()`: estimate the interested signatures enrolled in IOBR R package, which involves TME-associated, tumor-metabolism, and tumor-intrinsic signatures.
  - `feature_manipulation()`: manipulate features including the cell fraction and signatures generated from multi-omics data for latter analysis and model construction. Remove missing values, outliers and variables without significant variance.
  - `format_signatures()`: generate the object of `calculate_sig_score()`function, by inputting a data frame with signatures as column names of corresponding gene sets, and return a list contain the signature information for calculating multiple signature scores.
  - `format_msigdb()`: transform the signature gene sets data with gmt format, which is not included in the signature collection and might be downloaded in the MSgiDB website, into the object of `calculate_sig_score()`function.
  - **Batch Analysis and Visualization: batch survival analysis and batch correlation analysis and other batch statistical analyses**

    - `batch_surv`: batch survival analysis of multiple continuous variables including varied signature scores.
    - `subgroup_survival`: batch survival analysis of multiple categorized variables with different number of subgroups.
    - `batch_cor()`: batch analysis of correlation between two continuous variables using Pearson correlation coefficient or Spearman’s rank correlation coefficient .
    - `batch_wilcoxon()`: conduct batch wilcoxon analyses of binary variables.
    - `batch_pcc()`: batch analyses of Partial Correlation coefficient(PCC) between continuous variables and minimize the interference derived from confounding factors.
    - `iobr_cor_plot()`: visualization of batch correlation analysis of signatures from ‘sig\_group’. Visualize the correlation between signature or phenotype with expression of gene sets in target signature is also supported.
    - `cell_bar_plot()`: batch visualization of TME cell fraction, supporting input of deconvolution results from ‘CIBERSORT’, ‘EPIC’ and ‘quanTIseq’ methodologies to further compare the TME cell distributions within one sample or among different samples.
- **Signature Associated Mutation Module: identify and analyze mutations relevant to targeted signatures**

  - `make_mut_matrix()`: transform the mutation data with MAF format(contain the columns of gene ID and the corresponding gene alterations which including SNP, indel and frameshift) into a mutation matrix in a suitable manner for further investigating signature relevant mutations.
  - `find_mutations()`: identify mutations associated with a distinct phenotype or signature.
- **Model Construction Module: feature selection and fast model construct to predict clinical phenotype**

  - `BinomialModel()`: select features and construct a model to predict a binary phenotype.
  - `PrognosticMode()`: select features and construct a model to predict clinical survial outcome.

### 5. Functional Modules

IOBR consists of four functional modules, comprising decoding immune contexture (TME deconvolution module), estimation of signature scores, phenotype related signatures and corresponding genes, and signatures generated from single-cell sequencing data (signature module), analysis of signature associated mutations (signature associated mutation module) and fast model construction (model construction module). Details of each module and functional codes are described below. Hence, let us follow the analytic pipeline of IOBR step by step.

#### 5.1 Signature And TME Deconvolution Module

##### 5.2.1 Signature Estimation

IOBR integrates 255 published signature gene sets, involving tumor microenvironment, tumor metabolism, m6A, exosomes, microsatellite instability, and tertiary lymphoid structure. Running the function `signature_collection_citation` to attain the source papers. The function `signature_collection` returns the detail signature genes of all given signatures.

###### 1) Signature collection

Obtain the included signatures first.The signature collection is mainly classified into 3 categories: TME-associated, tumor-metabolism, and tumor-intrinsic signatures.

```
# Return available parameter options of signature estimation.
signature_score_calculation_methods
#>           PCA        ssGSEA       z-score   Integration 
#>         "pca"      "ssgsea"      "zscore" "integration"
#TME associated signatures
names(signature_tme)[1:20]
#>  [1] "CD_8_T_effector"            "DDR"                       
#>  [3] "APM"                        "Immune_Checkpoint"         
#>  [5] "CellCycle_Reg"              "Pan_F_TBRs"                
#>  [7] "Histones"                   "EMT1"                      
#>  [9] "EMT2"                       "EMT3"                      
#> [11] "WNT_target"                 "FGFR3_related"             
#> [13] "Cell_cycle"                 "Mismatch_Repair"           
#> [15] "Homologous_recombination"   "Nucleotide_excision_repair"
#> [17] "DNA_replication"            "Base_excision_repair"      
#> [19] "TMEscoreA_CIR"              "TMEscoreB_CIR"
#Metabolism related signatures
names(signature_metabolism)[1:20]
#>  [1] "Cardiolipin_Metabolism"                    
#>  [2] "Cardiolipin_Biosynthesis"                  
#>  [3] "Cholesterol_Biosynthesis"                  
#>  [4] "Citric_Acid_Cycle"                         
#>  [5] "Cyclooxygenase_Arachidonic_Acid_Metabolism"
#>  [6] "Prostaglandin_Biosynthesis"                
#>  [7] "Purine_Biosynthesis"                       
#>  [8] "Pyrimidine_Biosynthesis"                   
#>  [9] "Dopamine_Biosynthesis"                     
#> [10] "Epinephrine_Biosynthesis"                  
#> [11] "Norepinephrine_Biosynthesis"               
#> [12] "Fatty_Acid_Degradation"                    
#> [13] "Fatty_Acid_Elongation"                     
#> [14] "Fatty_Acid_Biosynthesis"                   
#> [15] "Folate_One_Carbon_Metabolism"              
#> [16] "Folate_biosynthesis"                       
#> [17] "Gluconeogenesis"                           
#> [18] "Glycolysis"                                
#> [19] "Glycogen_Biosynthesis"                     
#> [20] "Glycogen_Degradation"
#Signatures associated with biomedical basic research: such as m6A and exosomes
names(signature_tumor)
#>  [1] "Nature_metabolism_Hypoxia"                
#>  [2] "Winter_hypoxia_signature"                 
#>  [3] "Hu_hypoxia_signature"                     
#>  [4] "Molecular_Cancer_m6A"                     
#>  [5] "MT_exosome"                               
#>  [6] "SR_exosome"                               
#>  [7] "Positive_regulation_of_exosomal_secretion"
#>  [8] "Negative_regulation_of_exosomal_secretion"
#>  [9] "Exosomal_secretion"                       
#> [10] "Exosome_assembly"                         
#> [11] "Extracellular_vesicle_biogenesis"         
#> [12] "MC_Review_Exosome1"                       
#> [13] "MC_Review_Exosome2"                       
#> [14] "CMLS_Review_Exosome"                      
#> [15] "Ferroptosis"                              
#> [16] "EV_Cell_2020"
#signature collection including all aforementioned signatures 
names(signature_collection)[1:20]
#>  [1] "CD_8_T_effector"            "DDR"                       
#>  [3] "APM"                        "Immune_Checkpoint"         
#>  [5] "CellCycle_Reg"              "Pan_F_TBRs"                
#>  [7] "Histones"                   "EMT1"                      
#>  [9] "EMT2"                       "EMT3"                      
#> [11] "WNT_target"                 "FGFR3_related"             
#> [13] "Cell_cycle"                 "Mismatch_Repair"           
#> [15] "Homologous_recombination"   "Nucleotide_excision_repair"
#> [17] "DNA_replication"            "Base_excision_repair"      
#> [19] "TMEscoreA_CIR"              "TMEscoreB_CIR"
#citation of signatures
signature_collection_citation[1:20,]
#> # A tibble: 20 x 6
#>    Signatures    `Published year` Journal     Title              PMID  DOI      
#>    <chr>                    <dbl> <chr>       <chr>              <chr> <chr>    
#>  1 CD_8_T_effec…             2018 Nature      TGFβ attenuates t… 2944… 10.1038/…
#>  2 DDR                       2018 Nature      TGFβ attenuates t… 2944… 10.1038/…
#>  3 APM                       2018 Nature      TGFβ attenuates t… 2944… 10.1038/…
#>  4 Immune_Check…             2018 Nature      TGFβ attenuates t… 2944… 10.1038/…
#>  5 CellCycle_Reg             2018 Nature      TGFβ attenuates t… 2944… 10.1038/…
#>  6 Pan_F_TBRs                2018 Nature      TGFβ attenuates t… 2944… 10.1038/…
#>  7 Histones                  2018 Nature      TGFβ attenuates t… 2944… 10.1038/…
#>  8 EMT1                      2018 Nature      TGFβ attenuates t… 2944… 10.1038/…
#>  9 EMT2                      2018 Nature      TGFβ attenuates t… 2944… 10.1038/…
#> 10 EMT3                      2018 Nature      TGFβ attenuates t… 2944… 10.1038/…
#> 11 WNT_target                2018 Nature      TGFβ attenuates t… 2944… 10.1038/…
#> 12 FGFR3_related             2018 Nature      TGFβ attenuates t… 2944… 10.1038/…
#> 13 Cell_cycle                2018 Nature      TGFβ attenuates t… 2944… 10.1038/…
#> 14 Mismatch_Rep…             2018 Nature      TGFβ attenuates t… 2944… 10.1038/…
#> 15 Homologous_r…             2018 Nature      TGFβ attenuates t… 2944… 10.1038/…
#> 16 Nucleotide_e…             2018 Nature      TGFβ attenuates t… 2944… 10.1038/…
#> 17 DNA_replicat…             2018 Nature      TGFβ attenuates t… 2944… 10.1038/…
#> 18 Base_excisio…             2018 Nature      TGFβ attenuates t… 2944… 10.1038/…
#> 19 TMEscoreA_CIR             2019 Cancer Imm… Tumor Microenviro… 3084… 10.1158/…
#> 20 TMEscoreB_CIR             2019 Cancer Imm… Tumor Microenviro… 3084… 10.1158/…
```

###### 2) Methods for signature calculation

Three methodologies were adopted in the process of signature score evaluation, comprising Single-sample Gene Set Enrichment Analysis (ssGSEA), Principal component analysis (PCA), and Z-score.

###### Calculate TME associated signatures-(through PCA method).

```
data("eset_stad")
sig_tme<-calculate_sig_score(pdata           = NULL,
                             eset            = eset_stad,
                             signature       = signature_tme,
                             method          = "pca",
                             mini_gene_count = 2)
sig_tme[1:5, 1:10]
#> # A tibble: 5 x 10
#>   Index ID    CD_8_T_effector    DDR    APM Immune_Checkpoi… CellCycle_Reg
#>   <int> <chr>           <dbl>  <dbl>  <dbl>            <dbl>         <dbl>
#> 1     1 TCGA…           2.41   1.02   0.782           0.833         -0.463
#> 2     2 TCGA…          -0.581 -1.94   0.172           0.0794        -0.584
#> 3     3 TCGA…           1.09   0.735 -0.417           0.671         -0.247
#> 4     4 TCGA…           5.69   3.19   1.33            3.09          -0.553
#> 5     5 TCGA…           2.23   0.329  1.25            1.79           0.985
#> # … with 3 more variables: Pan_F_TBRs <dbl>, Histones <dbl>, EMT1 <dbl>
```

###### Estimate TME associated signatures-(through ssGSEA method).

```
sig_tme<-calculate_sig_score(pdata           = NULL,
                             eset            = eset_stad,
                             signature       = signature_tme,
                             method          = "ssgsea",
                             mini_gene_count = 5)
```

```
sig_tme[1:5, 1:10]
#> # A tibble: 5 x 10
#>   ID    Index CD_8_T_effector   DDR   APM Immune_Checkpoi… CellCycle_Reg
#>   <chr> <int>           <dbl> <dbl> <dbl>            <dbl>         <dbl>
#> 1 TCGA…     1           0.456 0.368 0.555            0.340         0.362
#> 2 TCGA…     2           0.334 0.343 0.537            0.278         0.419
#> 3 TCGA…     3           0.400 0.370 0.548            0.307         0.443
#> 4 TCGA…     4           0.539 0.394 0.573            0.453         0.442
#> 5 TCGA…     5           0.447 0.362 0.569            0.381         0.452
#> # … with 3 more variables: Pan_F_TBRs <dbl>, EMT1 <dbl>, EMT2 <dbl>
```

###### Evaluate metabolism related signatures.

```
sig_meta<-calculate_sig_score(pdata           = NULL,
                              eset            = eset_stad,
                              signature       = signature_metabolism,
                              method          = "pca",
                              mini_gene_count = 2)
sig_meta[1:5, 1:10]
#> # A tibble: 5 x 10
#>   Index ID    Cardiolipin_Met… Cardiolipin_Bio… Cholesterol_Bio…
#>   <int> <chr>            <dbl>            <dbl>            <dbl>
#> 1     1 TCGA…           -0.634          -0.0453            1.93 
#> 2     2 TCGA…            0.349          -0.156            -1.22 
#> 3     3 TCGA…            0.789           0.378             1.48 
#> 4     4 TCGA…           -0.963          -0.0409           -0.697
#> 5     5 TCGA…            0.311          -0.0895            0.353
#> # … with 5 more variables: Citric_Acid_Cycle <dbl>,
#> #   Cyclooxygenase_Arachidonic_Acid_Metabolism <dbl>,
#> #   Prostaglandin_Biosynthesis <dbl>, Purine_Biosynthesis <dbl>,
#> #   Pyrimidine_Biosynthesis <dbl>
```

###### Analyze all collected signature scores (integrating three methods: PCA, ssGSEA and z-score).

```
sig_res<-calculate_sig_score(pdata           = NULL,
                             eset            = eset_stad,
                             signature       = signature_collection,
                             method          = "integration",
                             mini_gene_count = 2)
```

```
sig_res[1:5,1:5]
#> # A tibble: 5 x 5
#>   ID           Index CD_8_T_effector_PCA DDR_PCA APM_PCA
#>   <chr>        <int>               <dbl>   <dbl>   <dbl>
#> 1 TCGA-B7-5818     1               2.41    1.02    0.782
#> 2 TCGA-BR-4187     2              -0.581  -1.94    0.172
#> 3 TCGA-BR-4201     3               1.09    0.735  -0.417
#> 4 TCGA-BR-4253     4               5.69    3.19    1.33 
#> 5 TCGA-BR-4256     5               2.23    0.329   1.25
```

###### The signature gene sets derived from GO, KEGG, HALLMARK and REACTOME datasets.

IOBR also enrolls the signature gene sets, containing GO, KEGG, HALLMARK and REACTOME gene sets obtained from MsigDB. The ssGSEA method is recommended for these signature estimation. This process may take a while for big datasets or calculating a large number of signatures.

```
sig_hallmark<-calculate_sig_score(pdata           = NULL,
                                  eset            = eset_stad,
                                  signature       = hallmark,
                                  method          = "ssgsea",
                                  mini_gene_count = 2)
```

```
sig_hallmark[1:5,1:5]
#> # A tibble: 5 x 5
#>   ID       Index HALLMARK_TNFA_SIGNALIN… HALLMARK_HYPOXIA HALLMARK_CHOLESTEROL_…
#>   <chr>    <int>                   <dbl>            <dbl>                  <dbl>
#> 1 TCGA-B7…     1                   0.911            0.794                  0.887
#> 2 TCGA-BR…     2                   0.923            0.865                  0.889
#> 3 TCGA-BR…     3                   0.980            0.870                  0.936
#> 4 TCGA-BR…     4                   0.952            0.754                  0.858
#> 5 TCGA-BR…     5                   1.01             0.863                  0.907
```

##### 5.2.2 Signatures Derived From Single Cell RNAseq

The recent advances in single-cell analysis make it a popular alternative to determine cell markers and gene signatures for phenotype. However, the significantly expensive cost and high requirement for starting tumor material still limited its widespread utility. Therefore, the attainable bulk sequencing larger dataset is still a need in validating the clinical significance of these signatures.

###### 1) Methods for signature estimation

Given that single-cell sequencing have a high resolution for dissecting signature genes of cells, we provide mutiple methodologies to extract cell signaturte genes from the single-cell sequencing data (`inputTPM` or `inputcounts`). Additionally, the linear `svr` algorithm of CIBERSORT or `lsei` algorithm is adopted in the bulk-seq data analysis to verify the clinical significance of the targeted cells identified by single-cell RNA sequencing.

Data Sources: Lineage-dependent gene expression programs influence the immune landscape of colorectal cancer. Nat Genet 2020 Method: 10X 3’ Data operation process: Cell filtering: 1. Only immune cells and stromal cells detected in the tumor are retained.

2. Select the cell type with the cell volume greater than 500.

Gene filtering: Genes detected in more than 5% of cells.

```
immune_feature_limma <- GenerateRef(dat     = inputTPM, 
                                    pheno   = pdatacounts$cluster,
                                    method  = "limma", 
                                    FDR     = 0.05)
#> NULL

immune_feature_DESeq2 <- GenerateRef(dds    = inputcounts,
                                     pheno  = pdatacounts$cluster, 
                                     dat    = inputTPM, 
                                     method = "DESeq2",
                                     FDR    = 0.05)
#> NULL

reference        <- immune_feature_DESeq2$reference_matrix
condition_number <- immune_feature_DESeq2$condition_number
top_probe        <- immune_feature_DESeq2$G
```

```
data("eset_stad")
svm<-deconvo_tme(eset            = eset_stad, 
                 reference       = reference, 
                 method          = "svm", 
                 arrays          = FALSE,
                 absolute.mode   = FALSE,
                 # abs.method    = "sig.score",
                 perm            = 1000)
head(svm)
#> # A tibble: 6 x 24
#>   ID    BcellsCD19CD20B… BcellsIgAPlasma… BcellsIgGPlasma… MyeloidsProinfl…
#>   <chr>            <dbl>            <dbl>            <dbl>            <dbl>
#> 1 TCGA…           0.188            0               0.00352                0
#> 2 TCGA…           0.0677           0               0.00207                0
#> 3 TCGA…           0.0768           0               0.0178                 0
#> 4 TCGA…           0.103            0.0270          0.0109                 0
#> 5 TCGA…           0.119            0               0.00757                0
#> 6 TCGA…           0.0875           0.0100          0                      0
#> # … with 19 more variables: MyeloidsProliferating_svm <dbl>,
#> #   MyeloidsSPP1_svm <dbl>, MyeloidscDC_svm <dbl>,
#> #   StromalcellsMyofibroblasts_svm <dbl>, StromalcellsPericytes_svm <dbl>,
#> #   StromalcellsStalklikeECs_svm <dbl>, StromalcellsStromal2_svm <dbl>,
#> #   StromalcellsStromal3_svm <dbl>, StromalcellsTiplikeECs_svm <dbl>,
#> #   TcellsCD4Tcells_svm <dbl>, TcellsCD8Tcells_svm <dbl>,
#> #   TcellsNKcells_svm <dbl>, TcellsRegulatoryTcells_svm <dbl>,
#> #   TcellsTfollicularhelpercells_svm <dbl>, TcellsThelper17cells_svm <dbl>,
#> #   TcellsgammadeltaTcells_svm <dbl>, `P-value_svm` <dbl>,
#> #   Correlation_svm <dbl>, RMSE_svm <dbl>

lsei<-deconvo_tme(eset           = eset_stad, 
                 reference       = reference, 
                 method          = "lsei", 
                 arrays          = FALSE,
                 scale_reference = T,
                 perm            = 1000)
head(lsei)
#> # A tibble: 6 x 21
#>   ID    BcellsCD19CD20B… BcellsIgAPlasma… BcellsIgGPlasma… MyeloidsProinfl…
#>   <chr>            <dbl>            <dbl>            <dbl>            <dbl>
#> 1 TCGA…                0                0          0.0181                 0
#> 2 TCGA…                0                0          0.0173                 0
#> 3 TCGA…                0                0          0.0324                 0
#> 4 TCGA…                0                0          0.0310                 0
#> 5 TCGA…                0                0          0.0241                 0
#> 6 TCGA…                0                0          0.00999                0
#> # … with 16 more variables: MyeloidsProliferating_lsei <dbl>,
#> #   MyeloidsSPP1_lsei <dbl>, MyeloidscDC_lsei <dbl>,
#> #   StromalcellsMyofibroblasts_lsei <dbl>, StromalcellsPericytes_lsei <dbl>,
#> #   StromalcellsStalklikeECs_lsei <dbl>, StromalcellsStromal2_lsei <dbl>,
#> #   StromalcellsStromal3_lsei <dbl>, StromalcellsTiplikeECs_lsei <dbl>,
#> #   TcellsCD4Tcells_lsei <dbl>, TcellsCD8Tcells_lsei <dbl>,
#> #   TcellsNKcells_lsei <dbl>, TcellsRegulatoryTcells_lsei <dbl>,
#> #   TcellsTfollicularhelpercells_lsei <dbl>, TcellsThelper17cells_lsei <dbl>,
#> #   TcellsgammadeltaTcells_lsei <dbl>
```

###### 2) Signature-phenotype correlation

Following is the example to explore the correlation between TLS score with tumor molecular characteristics, tumor microenvironment, and clinical phenotype of patients. B cells and tertiary lymphoid structures promote immunotherapy response

```
# Construct the signature list as an object first. The TLs signature mentioned above is included in the 'signature_collection' in IOBR.
signature_collection$TLS_Nature

# It is recommended to calculate other included signatures simultaneously, to systematically decode the correlation between TLS score and other tumor microenvironment characteristics.

# Data of mice received checkpoint blockade is adopted in this exploration. 
sig_res<-calculate_sig_score(pdata           = NULL,
                             eset            = eset_GSE63557,
                             signature       = signature_collection,
                             method          = "integration",
                             mini_gene_count = 2)
```

```
input<-merge(pdata_GSE63557,sig_res,by="ID",all =FALSE)
wilcox.test(input$TLS_Nature_PCA~input$BOR_binary)
#> 
#>  Wilcoxon rank sum test
#> 
#> data:  input$TLS_Nature_PCA by input$BOR_binary
#> W = 0, p-value = 1.083e-05
#> alternative hypothesis: true location shift is not equal to 0
```

##### 5.1.1 Available Methods to Decode TME Contexture

```
tme_deconvolution_methods
#>         MCPcounter               EPIC              xCell          CIBERSORT 
#>       "mcpcounter"             "epic"            "xcell"        "cibersort" 
#> CIBERSORT Absolute                IPS           ESTIMATE                SVM 
#>    "cibersort_abs"              "ips"         "estimate"              "svm" 
#>               lsei              TIMER          quanTIseq 
#>             "lsei"            "timer"        "quantiseq"
# Return available parameter options of deconvolution methods
```

If you use this package in your work, please cite both our package and the method(s) you are using.

##### Licenses of the deconvolution methods

| method | license | citation |
| --- | --- | --- |
| CIBERSORT | free for non-commerical use only | Newman, A. M., Liu, C. L., Green, M. R., Gentles, A. J., Feng, W., Xu, Y., … Alizadeh, A. A. (2015). Robust enumeration of cell subsets from tissue expression profiles. Nature Methods, 12(5), 453–457. https://doi.org/10.1038/nmeth.3337 |
| ESTIMATE | free (GPL2.0) | Vegesna R, Kim H, Torres-Garcia W, …, Verhaak R. (2013). Inferring tumour purity and stromal and immune cell admixture from expression data. Nature Communications 4, 2612. http://doi.org/10.1038/ncomms3612 |
| quanTIseq | free (BSD) | Finotello, F., Mayer, C., Plattner, C., Laschober, G., Rieder, D., Hackl, H., …, Sopper, S. (2019). Molecular and pharmacological modulators of the tumor immune contexture revealed by deconvolution of RNA-seq data. Genome medicine, 11(1), 34. https://doi.org/10.1186/s13073-019-0638-6 |
| TIMER | free (GPL 2.0) | Li, B., Severson, E., Pignon, J.-C., Zhao, H., Li, T., Novak, J., … Liu, X. S. (2016). Comprehensive analyses of tumor immunity: implications for cancer immunotherapy. Genome Biology, 17(1), 174. https://doi.org/10.1186/s13059-016-1028-7 |
| IPS | free (BSD) | P. Charoentong et al., Pan-cancer Immunogenomic Analyses Reveal Genotype-Immunophenotype Relationships and Predictors of Response to Checkpoint Blockade. Cell Reports 18, 248-262 (2017). https://doi.org/10.1016/j.celrep.2016.12.019 |
| MCPCounter | free (GPL 3.0) | Becht, E., Giraldo, N. A., Lacroix, L., Buttard, B., Elarouci, N., Petitprez, F., … de Reyniès, A. (2016). Estimating the population abundance of tissue-infiltrating immune and stromal cell populations using gene expression. Genome Biology, 17(1), 218. https://doi.org/10.1186/s13059-016-1070-5 |
| xCell | free (GPL 3.0) | Aran, D., Hu, Z., & Butte, A. J. (2017). xCell: digitally portraying the tissue cellular heterogeneity landscape. Genome Biology, 18(1), 220. https://doi.org/10.1186/s13059-017-1349-1 |
| EPIC | free for non-commercial use only (Academic License) | Racle, J., de Jonge, K., Baumgaertner, P., Speiser, D. E., & Gfeller, D. (2017). Simultaneous enumeration of cancer and immune cell types from bulk tumor gene expression data. ELife, 6, e26476. https://doi.org/10.7554/eLife.26476 |

##### Licenses of the signature-esitmation method

| method | license | citation |
| --- | --- | --- |
| GSVA | free (GPL (>= 2)) | Hänzelmann S, Castelo R, Guinney J (2013). “GSVA: gene set variation analysis for microarray and RNA-Seq data.” BMC Bioinformatics, 14, 7. doi: 10.1186/1471-2105-14-7, http://www.biomedcentral.com/1471-2105/14/7 |

The input data is a matrix (log2(TMP+1) transformed) containing 98 TCGA-STAD samples, with genes in rows and samples in columns. The row name must be HGNC symbols and the column name must be sample names.

```
# Load the `eset_stad` test data of gene expression matrix in IOBR
data("eset_stad")
eset_stad[1:5, 1:5]
#>         TCGA-B7-5818 TCGA-BR-4187 TCGA-BR-4201 TCGA-BR-4253 TCGA-BR-4256
#> MT-CO1      15.18012     15.55806     14.60960     14.63728     15.23528
#> MT-CO3      14.75536     15.19199     14.55337     13.54925     14.30425
#> MT-ND4      14.19637     15.61564     15.80262     14.98329     14.83764
#> MT-CO2      15.10790     15.50514     15.43261     14.52009     14.65806
#> MT-RNR2     14.22690     14.71157     13.48096     13.32553     13.55689
```

Check detail parameters of the function

```
help(deconvo_tme)
```

###### 1) Method 1: CIBERSORT

```
cibersort<-deconvo_tme(eset = eset_stad, method = "cibersort", arrays = FALSE, perm = 200 )
#> 
#> >>> Running CIBERSORT
head(cibersort)
#> # A tibble: 6 x 26
#>   ID    B_cells_naive_C… B_cells_memory_… Plasma_cells_CI… T_cells_CD8_CIB…
#>   <chr>            <dbl>            <dbl>            <dbl>            <dbl>
#> 1 TCGA…           0.0323                0          0                 0.193 
#> 2 TCGA…           0.0866                0          0                 0.0872
#> 3 TCGA…           0.0474                0          0.00644           0.0286
#> 4 TCGA…           0.0125                0          0.00257           0.224 
#> 5 TCGA…           0.0544                0          0.00923           0.0936
#> 6 TCGA…           0.0246                0          0.00162           0.124 
#> # … with 21 more variables: T_cells_CD4_naive_CIBERSORT <dbl>,
#> #   T_cells_CD4_memory_resting_CIBERSORT <dbl>,
#> #   T_cells_CD4_memory_activated_CIBERSORT <dbl>,
#> #   T_cells_follicular_helper_CIBERSORT <dbl>,
#> #   `T_cells_regulatory_(Tregs)_CIBERSORT` <dbl>,
#> #   T_cells_gamma_delta_CIBERSORT <dbl>, NK_cells_resting_CIBERSORT <dbl>,
#> #   NK_cells_activated_CIBERSORT <dbl>, Monocytes_CIBERSORT <dbl>,
#> #   Macrophages_M0_CIBERSORT <dbl>, Macrophages_M1_CIBERSORT <dbl>,
#> #   Macrophages_M2_CIBERSORT <dbl>, Dendritic_cells_resting_CIBERSORT <dbl>,
#> #   Dendritic_cells_activated_CIBERSORT <dbl>,
#> #   Mast_cells_resting_CIBERSORT <dbl>, Mast_cells_activated_CIBERSORT <dbl>,
#> #   Eosinophils_CIBERSORT <dbl>, Neutrophils_CIBERSORT <dbl>,
#> #   `P-value_CIBERSORT` <dbl>, Correlation_CIBERSORT <dbl>,
#> #   RMSE_CIBERSORT <dbl>
res<-cell_bar_plot(input = cibersort[1:12,], title = "CIBERSORT Cell Fraction")
#> There are seven categories you can choose: box, continue2, continue, random, heatmap, heatmap3, tidyheatmap
```

###### 2) Method 2: EPIC

```
help(deconvo_epic)
epic<-deconvo_tme(eset = eset_stad, method = "epic", arrays = FALSE)
#> 
#> >>> Running EPIC
#> Warning in EPIC::EPIC(bulk = eset, reference = ref, mRNA_cell = NULL, scaleExprs = TRUE): The optimization didn't fully converge for some samples:
#> TCGA-BR-6705; TCGA-BR-7958; TCGA-BR-8366; TCGA-BR-A4J4; TCGA-BR-A4J9; TCGA-CD-A4MG; TCGA-FP-8209; TCGA-HU-A4G9
#>  - check fit.gof for the convergeCode and convergeMessage
#> Warning in EPIC::EPIC(bulk = eset, reference = ref, mRNA_cell = NULL, scaleExprs
#> = TRUE): mRNA_cell value unknown for some cell types: CAFs, Endothelial - using
#> the default value of 0.4 for these but this might bias the true cell proportions
#> from all cell types.
head(epic)
#> # A tibble: 6 x 9
#>   ID    Bcells_EPIC CAFs_EPIC CD4_Tcells_EPIC CD8_Tcells_EPIC Endothelial_EPIC
#>   <chr>       <dbl>     <dbl>           <dbl>           <dbl>            <dbl>
#> 1 TCGA…      0.0226    0.0206           0.163          0.110             0.124
#> 2 TCGA…      0.0511    0.0254           0.189          0.107             0.182
#> 3 TCGA…      0.0435    0.0245           0.242          0.0726            0.146
#> 4 TCGA…      0.0353    0.0163           0.212          0.169             0.128
#> 5 TCGA…      0.0330    0.0226           0.208          0.111             0.177
#> 6 TCGA…      0.0145    0.0223           0.205          0.102             0.140
#> # … with 3 more variables: Macrophages_EPIC <dbl>, NKcells_EPIC <dbl>,
#> #   otherCells_EPIC <dbl>
```

###### 3) Method 3: MCPcounter

```
mcp<-deconvo_tme(eset = eset_stad, method = "mcpcounter")
#> 
#> >>> Running MCP-counter
head(mcp)
#> # A tibble: 6 x 11
#>   ID    T_cells_MCPcoun… CD8_T_cells_MCP… Cytotoxic_lymph… B_lineage_MCPco…
#>   <chr>            <dbl>            <dbl>            <dbl>            <dbl>
#> 1 TCGA…             2.26             2.81             1.59             1.62
#> 2 TCGA…             2.07             3.14             1.53             3.25
#> 3 TCGA…             2.28             1.11             1.96             3.46
#> 4 TCGA…             3.69             4.66             3.83             2.98
#> 5 TCGA…             2.71             3.40             2.26             3.01
#> 6 TCGA…             2.06             2.54             1.76             1.68
#> # … with 6 more variables: NK_cells_MCPcounter <dbl>,
#> #   Monocytic_lineage_MCPcounter <dbl>,
#> #   Myeloid_dendritic_cells_MCPcounter <dbl>, Neutrophils_MCPcounter <dbl>,
#> #   Endothelial_cells_MCPcounter <dbl>, Fibroblasts_MCPcounter <dbl>
```

###### 4) Method 4: xCELL

```
xcell<-deconvo_tme(eset = eset_stad, method = "xcell",arrays = FALSE)
```

```
head(xcell)
#> # A tibble: 6 x 68
#>   ID    Adipocytes_xCell Astrocytes_xCell `B-cells_xCell` Basophils_xCell
#>   <chr>            <dbl>            <dbl>           <dbl>           <dbl>
#> 1 TCGA…         0.                0.00159          0.0664        1.20e- 1
#> 2 TCGA…         8.96e- 3          0.132            0.0599        8.95e-19
#> 3 TCGA…         1.42e- 3          0.116            0.0504        6.64e-19
#> 4 TCGA…         4.12e-19          0                0.202         7.63e- 2
#> 5 TCGA…         7.68e- 3          0.127            0.0458        2.00e- 1
#> 6 TCGA…         1.34e-19          0.0508           0.0499        4.21e- 2
#> # … with 63 more variables: `CD4+_T-cells_xCell` <dbl>, `CD4+_Tcm_xCell` <dbl>,
#> #   `CD4+_Tem_xCell` <dbl>, `CD4+_memory_T-cells_xCell` <dbl>,
#> #   `CD4+_naive_T-cells_xCell` <dbl>, `CD8+_T-cells_xCell` <dbl>,
#> #   `CD8+_Tcm_xCell` <dbl>, `CD8+_Tem_xCell` <dbl>,
#> #   `CD8+_naive_T-cells_xCell` <dbl>, CLP_xCell <dbl>, CMP_xCell <dbl>,
#> #   Chondrocytes_xCell <dbl>, `Class-switched_memory_B-cells_xCell` <dbl>,
#> #   DC_xCell <dbl>, Endothelial_cells_xCell <dbl>, Eosinophils_xCell <dbl>,
#> #   Epithelial_cells_xCell <dbl>, Erythrocytes_xCell <dbl>,
#> #   Fibroblasts_xCell <dbl>, GMP_xCell <dbl>, HSC_xCell <dbl>,
#> #   Hepatocytes_xCell <dbl>, Keratinocytes_xCell <dbl>, MEP_xCell <dbl>,
#> #   MPP_xCell <dbl>, MSC_xCell <dbl>, Macrophages_xCell <dbl>,
#> #   Macrophages_M1_xCell <dbl>, Macrophages_M2_xCell <dbl>,
#> #   Mast_cells_xCell <dbl>, Megakaryocytes_xCell <dbl>,
#> #   Melanocytes_xCell <dbl>, `Memory_B-cells_xCell` <dbl>,
#> #   Mesangial_cells_xCell <dbl>, Monocytes_xCell <dbl>, Myocytes_xCell <dbl>,
#> #   NK_cells_xCell <dbl>, NKT_xCell <dbl>, Neurons_xCell <dbl>,
#> #   Neutrophils_xCell <dbl>, Osteoblast_xCell <dbl>, Pericytes_xCell <dbl>,
#> #   Plasma_cells_xCell <dbl>, Platelets_xCell <dbl>, Preadipocytes_xCell <dbl>,
#> #   Sebocytes_xCell <dbl>, Skeletal_muscle_xCell <dbl>,
#> #   Smooth_muscle_xCell <dbl>, Tgd_cells_xCell <dbl>, Th1_cells_xCell <dbl>,
#> #   Th2_cells_xCell <dbl>, Tregs_xCell <dbl>, aDC_xCell <dbl>, cDC_xCell <dbl>,
#> #   iDC_xCell <dbl>, ly_Endothelial_cells_xCell <dbl>,
#> #   mv_Endothelial_cells_xCell <dbl>, `naive_B-cells_xCell` <dbl>,
#> #   pDC_xCell <dbl>, `pro_B-cells_xCell` <dbl>, ImmuneScore_xCell <dbl>,
#> #   StromaScore_xCell <dbl>, MicroenvironmentScore_xCell <dbl>
```

###### 5) Method 5: ESTIMATE

```
estimate<-deconvo_tme(eset = eset_stad, method = "estimate")
#> [1] "Merged dataset includes 10156 genes (256 mismatched)."
#> [1] "1 gene set: StromalSignature  overlap= 139"
#> [1] "2 gene set: ImmuneSignature  overlap= 140"
head(estimate)
#> # A tibble: 6 x 5
#>   ID      StromalScore_est… ImmuneScore_esti… ESTIMATEScore_es… TumorPurity_est…
#>   <chr>               <dbl>             <dbl>             <dbl>            <dbl>
#> 1 TCGA-B…             -111.             1762.             1651.            0.662
#> 2 TCGA-B…             2054.             2208.             4262.            0.334
#> 3 TCGA-B…             1411.             1770.             3181.            0.479
#> 4 TCGA-B…              483.             2905.             3389.            0.451
#> 5 TCGA-B…             1659.             2541.             4200.            0.342
#> 6 TCGA-B…              831.             1722.             2553.            0.557
```

###### 6) Method 6: TIMER

```
timer<-deconvo_tme(eset = eset_stad, method = "timer", group_list = rep("stad",dim(eset_stad)[2]))
#> [1] "Outlier genes: FTL IGF2 IGHA1 IGHM IGKC IGKV4-1 LYZ MT-ATP6 MT-CO1 MT-CO2 MT-CO3 MT-CYB MT-ND1 MT-ND2 MT-ND3 MT-ND4 MT-ND4L MT-RNR1 MT-RNR2 MT-TP PGC"
#> Standardizing Data across genes
head(timer)
#> # A tibble: 6 x 7
#>   ID    B_cell_TIMER T_cell_CD4_TIMER T_cell_CD8_TIMER Neutrophil_TIMER
#>   <chr>        <dbl>            <dbl>            <dbl>            <dbl>
#> 1 TCGA…       0.0954            0.130            0.191            0.114
#> 2 TCGA…       0.0983            0.131            0.202            0.125
#> 3 TCGA…       0.0996            0.122            0.200            0.127
#> 4 TCGA…       0.101             0.131            0.240            0.129
#> 5 TCGA…       0.0945            0.133            0.213            0.137
#> 6 TCGA…       0.0907            0.126            0.199            0.121
#> # … with 2 more variables: Macrophage_TIMER <dbl>, DC_TIMER <dbl>
```

###### 7) Method 7: quanTIseq

```
quantiseq<-deconvo_tme(eset = eset_stad, tumor = TRUE, arrays = FALSE, scale_mrna = TRUE, method = "quantiseq")
#> 
#> Running quanTIseq deconvolution module
#> Gene expression normalization and re-annotation (arrays: FALSE)
#> Removing 17 noisy genes
#> Removing 15 genes with high expression in tumors
#> Signature genes found in data set: 138/138 (100%)
#> Mixture deconvolution (method: lsei)
#> Deconvolution sucessful!
head(quantiseq)
#> # A tibble: 6 x 12
#>   ID    B_cells_quantis… Macrophages_M1_… Macrophages_M2_… Monocytes_quant…
#>   <chr>            <dbl>            <dbl>            <dbl>            <dbl>
#> 1 TCGA…          0.00897           0.0943           0.0299                0
#> 2 TCGA…          0.0269            0.0155           0.169                 0
#> 3 TCGA…          0.0160            0.0570           0.0661                0
#> 4 TCGA…          0.0240            0.171            0.128                 0
#> 5 TCGA…          0.0129            0.126            0.0922                0
#> 6 TCGA…          0.0148            0.0685           0.0673                0
#> # … with 7 more variables: Neutrophils_quantiseq <dbl>,
#> #   NK_cells_quantiseq <dbl>, T_cells_CD4_quantiseq <dbl>,
#> #   T_cells_CD8_quantiseq <dbl>, Tregs_quantiseq <dbl>,
#> #   Dendritic_cells_quantiseq <dbl>, Other_quantiseq <dbl>
res<-cell_bar_plot(input = quantiseq[1:12, ], title = "quanTIseq Cell Fraction")
#> There are seven categories you can choose: box, continue2, continue, random, heatmap, heatmap3, tidyheatmap
```

###### 8) Method 8: IPS

```
ips<-deconvo_tme(eset = eset_stad, method = "ips", plot= FALSE)
head(ips)
#> # A tibble: 6 x 7
#>   ID           MHC_IPS EC_IPS SC_IPS CP_IPS AZ_IPS IPS_IPS
#>   <chr>          <dbl>  <dbl>  <dbl>  <dbl>  <dbl>   <dbl>
#> 1 TCGA-B7-5818    3.56   1.09  -1.37 -0.576   2.70       9
#> 2 TCGA-BR-4187    3.37   1.18  -1.71 -0.308   2.53       8
#> 3 TCGA-BR-4201    3.04   1.22  -1.62 -0.439   2.20       7
#> 4 TCGA-BR-4253    3.79   1.63  -1.74 -1.25    2.42       8
#> 5 TCGA-BR-4256    3.65   1.40  -1.95 -0.847   2.25       8
#> 6 TCGA-BR-4257    3.11   1.23  -1.74 -0.529   2.07       7
```

###### 9) Combination of above deconvolution results

```
tme_combine<-cibersort %>% 
  inner_join(.,mcp,by       = "ID") %>% 
  inner_join(.,xcell,by     = "ID") %>%
  inner_join(.,epic,by      = "ID") %>% 
  inner_join(.,estimate,by  = "ID") %>% 
  inner_join(.,timer,by     = "ID") %>% 
  inner_join(.,quantiseq,by = "ID") %>% 
  inner_join(.,ips,by       = "ID")
dim(tme_combine)
#> [1]  98 138
```

#### 5.2 Phenotype Module

##### 5.2.3 Explore Phenotype Relevant Signatures

###### 1) Load data

Data of IMvigor210 immunotherapy cohort was used for batch analysis of phenotype-related signatures.

```
## Load the signatures and Cell fractions calculated in previous steps by IOBR.
data("imvigor210_sig")
imvigor210_sig[1:5, 1:10]
#> # A tibble: 5 x 10
#>   ID    B_cells_naive_C… B_cells_memory_… Plasma_cells_CI… T_cells_CD8_CIB…
#>   <chr>            <dbl>            <dbl>            <dbl>            <dbl>
#> 1 SAMf…           0.0290          0.0457           0                      0
#> 2 SAM6…           0.0812          0.00127          0                      0
#> 3 SAMc…           0.0124          0                0.00127                0
#> 4 SAM8…           0               0                0.00119                0
#> 5 SAMf…           0               0                0.00949                0
#> # … with 5 more variables: T_cells_CD4_naive_CIBERSORT <dbl>,
#> #   T_cells_CD4_memory_resting_CIBERSORT <dbl>,
#> #   T_cells_CD4_memory_activated_CIBERSORT <dbl>,
#> #   T_cells_follicular_helper_CIBERSORT <dbl>,
#> #   `T_cells_regulatory_(Tregs)_CIBERSORT` <dbl>

# Check therapeutic response and survival outcome in the phenotype data.
data("imvigor210_pdata")
imvigor210_pdata[1:5, 1:5]
#> # A tibble: 5 x 5
#>   ID              BOR   BOR_binary OS_days            OS_status
#>   <chr>           <chr> <chr>      <chr>              <chr>    
#> 1 SAM00b9e5c52da9 NA    NA         57.166324439999997 1        
#> 2 SAM0257bbbbd388 SD    NR         469.15811100000002 1        
#> 3 SAM025b45c27e05 PD    NR         263.16221766000001 1        
#> 4 SAM032c642382a7 PD    NR         74.907597539999998 1        
#> 5 SAM04c589eb3fb3 NA    NA         20.698151939999999 0
```

###### 2)Signature groups

Run `sig_group()` function for latter batch analysis of response associated signatures.

```
# The signature group list.
names(sig_group)
#>  [1] "tumor_signature"        "EMT"                    "io_biomarkers"         
#>  [4] "immu_microenvironment"  "immu_suppression"       "immu_exclusion"        
#>  [7] "immu_exhaustion"        "TCR_BCR"                "tme_signatures1"       
#> [10] "tme_signatures2"        "Bcells"                 "Tcells"                
#> [13] "DCs"                    "Macrophages"            "Neutrophils"           
#> [16] "Monocytes"              "CAFs"                   "NK"                    
#> [19] "tme_cell_types"         "CIBERSORT"              "MCPcounter"            
#> [22] "EPIC"                   "xCell"                  "quanTIseq"             
#> [25] "ESTIMATE"               "IPS"                    "TIMER"                 
#> [28] "fatty_acid_metabolism"  "hypoxia_signature"      "cholesterol_metabolism"
#> [31] "Metabolism"             "hallmark"               "hallmark1"             
#> [34] "hallmark2"              "hallmark3"              "Rooney_et_al"          
#> [37] "Bindea_et_al"           "Li_et_al"               "Peng_et_al"

# The tumor-relevant signatures in first group.
names(sig_group)[1]
#> [1] "tumor_signature"
sig_group[[1]]
#>  [1] "CellCycle_Reg"                            
#>  [2] "Cell_cycle"                               
#>  [3] "DDR"                                      
#>  [4] "Mismatch_Repair"                          
#>  [5] "Histones"                                 
#>  [6] "Homologous_recombination"                 
#>  [7] "Nature_metabolism_Hypoxia"                
#>  [8] "Molecular_Cancer_m6A"                     
#>  [9] "MT_exosome"                               
#> [10] "Positive_regulation_of_exosomal_secretion"
#> [11] "Ferroptosis"                              
#> [12] "EV_Cell_2020"

# The signatures associated with immunotherapy biomarkers.
names(sig_group)[3]
#> [1] "io_biomarkers"
sig_group[[3]]
#>  [1] "TMEscore_CIR"                    "TMEscoreA_CIR"                  
#>  [3] "TMEscoreB_CIR"                   "T_cell_inflamed_GEP_Ayers_et_al"
#>  [5] "CD_8_T_effector"                 "IPS_IPS"                        
#>  [7] "Immune_Checkpoint"               "Exhausted_CD8_Danaher_et_al"    
#>  [9] "Pan_F_TBRs"                      "Mismatch_Repair"                
#> [11] "APM"

# The signatures of immunosuppression.
names(sig_group)[5]
#> [1] "immu_suppression"
sig_group[[5]]
#> [1] "Pan_F_TBRs"                          
#> [2] "Fibroblasts_MCPcounter"              
#> [3] "Immune_Checkpoint"                   
#> [4] "Exhausted_CD8_Danaher_et_al"         
#> [5] "MDSC_Wang_et_al"                     
#> [6] "Macrophages_M2_cibersort"            
#> [7] "Tregs_quantiseq"                     
#> [8] "T_cells_regulatory_(Tregs)_CIBERSORT"
```

###### 3) Identify phenotype relevant signatures

Construct phenotype group and bath visualize correlation between therapy response and selected signatures.

```
pdata_group<-imvigor210_pdata[!imvigor210_pdata$BOR_binary=="NA",c("ID","BOR","BOR_binary")]
res<-iobr_cor_plot(pdata_group           = pdata_group,
                   id1                   = "ID",
                   feature_data          = imvigor210_sig,
                   id2                   = "ID",
                   target                = NULL,
                   group                 = "BOR_binary",
                   is_target_continuous  = FALSE,
                   padj_cutoff           = 1,
                   index                 = 1,
                   category              = "signature",
                   signature_group       = sig_group[c(1,3,5)],
                   ProjectID             = "IMvigor210",
                   palette_box           = "paired1",
                   palette_corplot       = "pheatmap",
                   palette_heatmap       = 2,
                   feature_limit         = 26,
                   character_limit       = 30,
                   show_heatmap_col_name = FALSE,
                   show_col              = FALSE,
                   show_plot             = TRUE)
#> [1] ">>>  Processing signature: tumor_signature"
```

```
#> [1] ">>>  Processing signature: io_biomarkers"
```

```
#> [1] ">>>  Processing signature: immu_suppression"
```

```
#> [1] ">>> Proportion of two groups:"
#>  NR   R 
#> 230  68
head(res)
#> # A tibble: 6 x 8
#>   sig_names             p.value     NR     R statistic   p.adj log10pvalue stars
#>   <chr>                   <dbl>  <dbl> <dbl>     <dbl>   <dbl>       <dbl> <fct>
#> 1 Mismatch_Repair       9.21e-6 -0.142 0.479    -0.620 1.22e-4        5.04 **** 
#> 2 Cell_cycle            1.05e-5 -0.143 0.484    -0.627 1.22e-4        4.98 **** 
#> 3 DDR                   1.52e-5 -0.140 0.474    -0.614 1.22e-4        4.82 **** 
#> 4 Homologous_recombi…   3.31e-5 -0.134 0.453    -0.587 1.79e-4        4.48 **** 
#> 5 Histones              3.72e-5 -0.127 0.429    -0.556 1.79e-4        4.43 **** 
#> 6 CD_8_T_effector       9.11e-4 -0.104 0.351    -0.455 3.64e-3        3.04 ***
```

###### 4) Visualize phenotype relevant signature genes

Estimate and visualize the expression of response relevant signature genes.

```
imvigor210_eset[1:5, 1:6]
#>              SAMf2ce197162ce SAM698d8d76b934 SAMc1b27bc16435 SAM85e41e7f33f9
#> LOC100093631       3.1009398       2.8207636       3.7058993      2.81012848
#> LOC100126784      -2.6237747      -4.2560520      -5.4104447     -2.07600356
#> LOC100128108      -1.5017841       0.5200520      -0.7665885     -2.07600356
#> LOC100128288      -0.3361981      -1.2204281      -1.9510131     -1.25886761
#> LOC100128361       0.2545468       0.2923847      -0.2009913     -0.02537748
#>              SAMf275eb859a39 SAM7f0d9cc7f001
#> LOC100093631       4.0102463      5.24697867
#> LOC100126784      -3.6376118     -2.23243417
#> LOC100128108      -1.9495558      0.08949393
#> LOC100128288       0.3320146     -0.33431378
#> LOC100128361       0.7698920     -1.16830383

res<-iobr_cor_plot(pdata_group           = pdata_group,
                   id1                   = "ID",
                   feature_data          = imvigor210_eset,
                   id2                   = "ID",
                   target                = NULL,
                   group                 = "BOR_binary",
                   is_target_continuous  = FALSE,
                   padj_cutoff           = 1,
                   index                 = 2,
                   category              = "gene",
                   signature_group       = signature_collection[c(1:2,4)],    
                   ProjectID             = "IMvigor210",
                   palette_box           = "paired1",
                   palette_corplot       = "pheatmap",
                   palette_heatmap       = 4,
                   feature_limit         = 26,
                   character_limit       = 30,
                   show_heatmap_col_name = FALSE,
                   show_col              = FALSE,
                   show_plot             = TRUE)
#> [1] ">>>  Processing signature: CD_8_T_effector"
```

```
#> [1] ">>>  Processing signature: DDR"
```

```
#> [1] ">>>  Processing signature: Immune_Checkpoint"
```

```
#> [1] ">>> Proportion of two groups:"
#>  NR   R 
#> 230  68
head(res)
#> # A tibble: 6 x 8
#>   sig_names    p.value     NR     R statistic    p.adj log10pvalue stars
#>   <chr>          <dbl>  <dbl> <dbl>     <dbl>    <dbl>       <dbl> <fct>
#> 1 RDM1      0.00000113 -0.147 0.499    -0.646 0.000205        5.95 **** 
#> 2 ERCC4     0.00000301 -0.154 0.522    -0.677 0.000272        5.52 **** 
#> 3 FANCI     0.00000643 -0.140 0.473    -0.612 0.000388        5.19 **** 
#> 4 EME1      0.0000173  -0.132 0.446    -0.578 0.000783        4.76 **** 
#> 5 POLE      0.0000246  -0.133 0.451    -0.584 0.000890        4.61 **** 
#> 6 BLM       0.0000295  -0.130 0.441    -0.571 0.000891        4.53 ****
```

###### 5) Estimate lncRNA associated signatures

```
## Load the signatures and Cell fractions calculated in previous steps by IOBR.
head(as_tibble(imvigor210_sig))
#> # A tibble: 6 x 456
#>   ID    B_cells_naive_C… B_cells_memory_… Plasma_cells_CI… T_cells_CD8_CIB…
#>   <chr>            <dbl>            <dbl>            <dbl>            <dbl>
#> 1 SAMf…           0.0290          0.0457           0                      0
#> 2 SAM6…           0.0812          0.00127          0                      0
#> 3 SAMc…           0.0124          0                0.00127                0
#> 4 SAM8…           0               0                0.00119                0
#> 5 SAMf…           0               0                0.00949                0
#> 6 SAM7…           0.0582          0.129            0                      0
#> # … with 451 more variables: T_cells_CD4_naive_CIBERSORT <dbl>,
#> #   T_cells_CD4_memory_resting_CIBERSORT <dbl>,
#> #   T_cells_CD4_memory_activated_CIBERSORT <dbl>,
#> #   T_cells_follicular_helper_CIBERSORT <dbl>,
#> #   `T_cells_regulatory_(Tregs)_CIBERSORT` <dbl>,
#> #   T_cells_gamma_delta_CIBERSORT <dbl>, NK_cells_resting_CIBERSORT <dbl>,
#> #   NK_cells_activated_CIBERSORT <dbl>, Monocytes_CIBERSORT <dbl>,
#> #   Macrophages_M0_CIBERSORT <dbl>, Macrophages_M1_CIBERSORT <dbl>,
#> #   Macrophages_M2_CIBERSORT <dbl>, Dendritic_cells_resting_CIBERSORT <dbl>,
#> #   Dendritic_cells_activated_CIBERSORT <dbl>,
#> #   Mast_cells_resting_CIBERSORT <dbl>, Mast_cells_activated_CIBERSORT <dbl>,
#> #   Eosinophils_CIBERSORT <dbl>, Neutrophils_CIBERSORT <dbl>,
#> #   T_cells_MCPcounter <dbl>, CD8_T_cells_MCPcounter <dbl>,
#> #   Cytotoxic_lymphocytes_MCPcounter <dbl>, NK_cells_MCPcounter <dbl>,
#> #   B_lineage_MCPcounter <dbl>, Monocytic_lineage_MCPcounter <dbl>,
#> #   Myeloid_dendritic_cells_MCPcounter <dbl>, Neutrophils_MCPcounter <dbl>,
#> #   Endothelial_cells_MCPcounter <dbl>, Fibroblasts_MCPcounter <dbl>,
#> #   aDC_xCell <dbl>, Adipocytes_xCell <dbl>, Astrocytes_xCell <dbl>,
#> #   `B-cells_xCell` <dbl>, Basophils_xCell <dbl>,
#> #   `CD4+_memory_T-cells_xCell` <dbl>, `CD4+_naive_T-cells_xCell` <dbl>,
#> #   `CD4+_T-cells_xCell` <dbl>, `CD4+_Tcm_xCell` <dbl>, `CD4+_Tem_xCell` <dbl>,
#> #   `CD8+_naive_T-cells_xCell` <dbl>, `CD8+_T-cells_xCell` <dbl>,
#> #   `CD8+_Tcm_xCell` <dbl>, `CD8+_Tem_xCell` <dbl>, cDC_xCell <dbl>,
#> #   Chondrocytes_xCell <dbl>, `Class-switched_memory_B-cells_xCell` <dbl>,
#> #   CLP_xCell <dbl>, CMP_xCell <dbl>, DC_xCell <dbl>,
#> #   Endothelial_cells_xCell <dbl>, Eosinophils_xCell <dbl>,
#> #   Epithelial_cells_xCell <dbl>, Erythrocytes_xCell <dbl>,
#> #   Fibroblasts_xCell <dbl>, GMP_xCell <dbl>, Hepatocytes_xCell <dbl>,
#> #   HSC_xCell <dbl>, iDC_xCell <dbl>, Keratinocytes_xCell <dbl>,
#> #   ly_Endothelial_cells_xCell <dbl>, Macrophages_xCell <dbl>,
#> #   Macrophages_M1_xCell <dbl>, Macrophages_M2_xCell <dbl>,
#> #   Mast_cells_xCell <dbl>, Megakaryocytes_xCell <dbl>,
#> #   Melanocytes_xCell <dbl>, `Memory_B-cells_xCell` <dbl>, MEP_xCell <dbl>,
#> #   Mesangial_cells_xCell <dbl>, Monocytes_xCell <dbl>, MPP_xCell <dbl>,
#> #   MSC_xCell <dbl>, mv_Endothelial_cells_xCell <dbl>, Myocytes_xCell <dbl>,
#> #   `naive_B-cells_xCell` <dbl>, Neurons_xCell <dbl>, Neutrophils_xCell <dbl>,
#> #   NK_cells_xCell <dbl>, NKT_xCell <dbl>, Osteoblast_xCell <dbl>,
#> #   pDC_xCell <dbl>, Pericytes_xCell <dbl>, Plasma_cells_xCell <dbl>,
#> #   Platelets_xCell <dbl>, Preadipocytes_xCell <dbl>,
#> #   `pro_B-cells_xCell` <dbl>, Sebocytes_xCell <dbl>,
#> #   Skeletal_muscle_xCell <dbl>, Smooth_muscle_xCell <dbl>,
#> #   Tgd_cells_xCell <dbl>, Th1_cells_xCell <dbl>, Th2_cells_xCell <dbl>,
#> #   Tregs_xCell <dbl>, ImmuneScore_xCell <dbl>, StromaScore_xCell <dbl>,
#> #   MicroenvironmentScore_xCell <dbl>, Bcells_EPIC <dbl>, CAFs_EPIC <dbl>,
#> #   CD4_Tcells_EPIC <dbl>, CD8_Tcells_EPIC <dbl>, Endothelial_EPIC <dbl>, …

# Check therapeutic response and survival outcome in the phenotype data.
head(as_tibble(imvigor210_pdata))
#> # A tibble: 6 x 12
#>   ID    BOR   BOR_binary OS_days OS_status Mutation_Load Neo_antigen_Load
#>   <chr> <chr> <chr>      <chr>   <chr>     <chr>         <chr>           
#> 1 SAM0… NA    NA         57.166… 1         NA            NA              
#> 2 SAM0… SD    NR         469.15… 1         18            4.6862745099999…
#> 3 SAM0… PD    NR         263.16… 1         1             0.31372549      
#> 4 SAM0… PD    NR         74.907… 1         44            6.1960784310000…
#> 5 SAM0… NA    NA         20.698… 0         50            NA              
#> 6 SAM0… SD    NR         136.01… 1         2             1.4705882349999…
#> # … with 5 more variables: CD_8_T_effector <dbl>, Immune_Checkpoint <dbl>,
#> #   Pan_F_TBRs <chr>, Mismatch_Repair <chr>, TumorPurity <dbl>
```

###### Define the latent molecur functions of non-coding RNA by exploring lncRNA related signatures.

- Multiple literatures have reported that *HCP5* is correlated with tumor microenvironment and immunotherapeutic responses.Citation-1,Citation-2,Citation-3;
- Reports also suggested the association between *LINC00657* and progression of multiple cancers.Citation-1,Citation-2，Citation-3
- Taken *HCP5* and *LINC00657* for example, we explore their relationships with TME and other well-recognized signatures, to better reveal their potential functions.

```
# Load the test data: the gene expression matrix of IMvigor210 cohort has been normalized using method `voom`.
imvigor210_eset[1:5,1:5]
#>              SAMf2ce197162ce SAM698d8d76b934 SAMc1b27bc16435 SAM85e41e7f33f9
#> LOC100093631       3.1009398       2.8207636       3.7058993      2.81012848
#> LOC100126784      -2.6237747      -4.2560520      -5.4104447     -2.07600356
#> LOC100128108      -1.5017841       0.5200520      -0.7665885     -2.07600356
#> LOC100128288      -0.3361981      -1.2204281      -1.9510131     -1.25886761
#> LOC100128361       0.2545468       0.2923847      -0.2009913     -0.02537748
#>              SAMf275eb859a39
#> LOC100093631       4.0102463
#> LOC100126784      -3.6376118
#> LOC100128108      -1.9495558
#> LOC100128288       0.3320146
#> LOC100128361       0.7698920
# Extract the significant genes as a phenotype of the patients based on the exploration of expression matrix.
pdata_group<-rownames_to_column(as.data.frame(t(imvigor210_eset)),var = "ID")

pdata_group<-as.data.frame(pdata_group[,c("ID","HCP5","LINC00657")])
head(pdata_group)
#>                ID       HCP5 LINC00657
#> 1 SAMf2ce197162ce -1.8532565  2.746143
#> 2 SAM698d8d76b934 -1.7199991  1.636339
#> 3 SAMc1b27bc16435 -2.6030898  3.863351
#> 4 SAM85e41e7f33f9 -0.5375836  2.989060
#> 5 SAMf275eb859a39  0.4685876  2.599731
#> 6 SAM7f0d9cc7f001 -1.4439383  2.029233
```

Utilize `iobr_cor_plot`function for batch statistical analysis and visualization of the correlation between *HCP5* and TME.

```
res<-iobr_cor_plot(pdata_group           = pdata_group,
                   id1                   = "ID",
                   feature_data          = imvigor210_sig,
                   id2                   = "ID",
                   target                = "HCP5",
                   group                 = "group3",
                   is_target_continuous  = TRUE,
                   padj_cutoff           = 1,
                   index                 = 3,
                   category              = "signature",
                   signature_group       = sig_group[1:3],
                   ProjectID             = "IMvigor210",
                   palette_box           = "set2",
                   palette_corplot       = "pheatmap",
                   palette_heatmap       = 2,
                   feature_limit         = 26,
                   character_limit       = 30,
                   show_heatmap_col_name = FALSE,
                   show_col              = FALSE,
                   show_plot             = TRUE)
#> [1] ">>>  Processing signature: tumor_signature"
```

```
#> [1] ">>>  Processing signature: EMT"
```

```
#> [1] ">>>  Processing signature: io_biomarkers"
```

```
# Check targeted genes and the statistical correlations with all analyzed signatures. (determined by Spearman's rank correlation coefficient.)  
head(res)
#> # A tibble: 6 x 6
#>   sig_names                        p.value statistic    p.adj log10pvalue stars
#>   <chr>                              <dbl>     <dbl>    <dbl>       <dbl> <fct>
#> 1 TMEscoreA_CIR                   1.44e-54     0.710 3.45e-53        53.8 **** 
#> 2 T_cell_inflamed_GEP_Ayers_et_al 3.09e-51     0.694 3.71e-50        50.5 **** 
#> 3 APM                             2.58e-49     0.684 2.07e-48        48.6 **** 
#> 4 Immune_Checkpoint               1.07e-46     0.670 6.44e-46        46.0 **** 
#> 5 CD_8_T_effector                 3.38e-44     0.656 1.62e-43        43.5 **** 
#> 6 Exhausted_CD8_Danaher_et_al     6.81e-36     0.603 2.72e-35        35.2 ****
```

Utilize `iobr_cor_plot`function for batch statistical analysis and visualization of the correlation between *LINC00657* and TME.

```
# Only part of the sign_group was exhibited, as an example of the more extensive results.
names(sig_group)
#>  [1] "tumor_signature"        "EMT"                    "io_biomarkers"         
#>  [4] "immu_microenvironment"  "immu_suppression"       "immu_exclusion"        
#>  [7] "immu_exhaustion"        "TCR_BCR"                "tme_signatures1"       
#> [10] "tme_signatures2"        "Bcells"                 "Tcells"                
#> [13] "DCs"                    "Macrophages"            "Neutrophils"           
#> [16] "Monocytes"              "CAFs"                   "NK"                    
#> [19] "tme_cell_types"         "CIBERSORT"              "MCPcounter"            
#> [22] "EPIC"                   "xCell"                  "quanTIseq"             
#> [25] "ESTIMATE"               "IPS"                    "TIMER"                 
#> [28] "fatty_acid_metabolism"  "hypoxia_signature"      "cholesterol_metabolism"
#> [31] "Metabolism"             "hallmark"               "hallmark1"             
#> [34] "hallmark2"              "hallmark3"              "Rooney_et_al"          
#> [37] "Bindea_et_al"           "Li_et_al"               "Peng_et_al"
res<-iobr_cor_plot(pdata_group           = pdata_group,
                   id1                   = "ID",
                   feature_data          = imvigor210_sig,
                   id2                   = "ID",
                   target                = "LINC00657",
                   group                 = "group3",
                   is_target_continuous  = TRUE,
                   padj_cutoff           = 1,
                   index                 = 4,
                   category              = "signature",
                   signature_group       = sig_group[c(1,3:4)],
                   ProjectID             = "IMvigor210",
                   palette_box           = "jco",
                   palette_corplot       = "pheatmap",
                   palette_heatmap       = 4,
                   feature_limit         = 26,
                   character_limit       = 30,
                   show_heatmap_col_name = FALSE,
                   show_col              = FALSE,
                   show_plot             = TRUE)
#> [1] ">>>  Processing signature: tumor_signature"
```

```
#> [1] ">>>  Processing signature: io_biomarkers"
```

```
#> [1] ">>>  Processing signature: immu_microenvironment"
```

```
# # Check targeted genes and the statistical correlations with all analyzed signatures. (determined by Spearman's rank correlation coefficient.)  
head(res)
#> # A tibble: 6 x 6
#>   sig_names                         p.value statistic    p.adj log10pvalue stars
#>   <chr>                               <dbl>     <dbl>    <dbl>       <dbl> <fct>
#> 1 Positive_regulation_of_exosomal… 5.21e-19     0.453 1.46e-17        18.3 **** 
#> 2 TMEscoreA_CIR                    1.37e-16    -0.424 1.92e-15        15.9 **** 
#> 3 T_cell_inflamed_GEP_Ayers_et_al  2.94e-16    -0.419 2.74e-15        15.5 **** 
#> 4 CD_8_T_effector                  4.08e-15    -0.404 2.86e-14        14.4 **** 
#> 5 TAM_Peng_et_al                   7.10e-15    -0.401 3.36e-14        14.1 **** 
#> 6 Immune_Checkpoint                7.19e-15    -0.401 3.36e-14        14.1 ****
```

##### 5.2.4 Identify Signatures Relevant to Targeted Signature

###### 1) Construct phenotype group

Construct the `pdata_group`as a phenotype data frame of patients.

```
# The data have been saved in `imvigor210_pdata`.
# head(imvigor210_pdata)
pdata_group<-as.data.frame(imvigor210_pdata[,c("ID","Pan_F_TBRs")])
pdata_group$Pan_F_TBRs<-scale(as.numeric(pdata_group$Pan_F_TBRs))
head(pdata_group)
#>                ID Pan_F_TBRs
#> 1 SAM00b9e5c52da9  0.2681507
#> 2 SAM0257bbbbd388  0.4618378
#> 3 SAM025b45c27e05 -0.8229645
#> 4 SAM032c642382a7 -0.4156874
#> 5 SAM04c589eb3fb3  9.0208975
#> 6 SAM0571f17f4045 -0.4462966
```

###### 2) Analyze Pan-F-TBRs correlated signatures

`Pan-F-TBRs` is an immune-exclusion signature published in Nature in 2018: S. Mariathasan et al., TGFbeta attenuates tumour response to PD-L1 blockade by contributing to exclusion of T cells. Nature 554, 544-548 (2018).

The Pan-F-TBRs score obtained could be used to batch analyze and visualize the interaction between this immune-exclusion signature and other interested `signatures` , as well as its significant `cell fractions`, and to further explore other biomarkers may impact corresponding biological process.

```
res<-iobr_cor_plot(pdata_group           = pdata_group,
                   id1                   = "ID",
                   feature_data          = imvigor210_sig,
                   id2                   = "ID",
                   target                = "Pan_F_TBRs",
                   group                 = "group3",
                   is_target_continuous  = TRUE,
                   padj_cutoff           = 1,
                   index                 = 5,
                   category              = "signature",
                   signature_group       =sig_group[1:2],
                   ProjectID             = "IMvigor210",
                   palette_box           = "set2",
                   palette_corplot       = "pheatmap",
                   palette_heatmap       = 2,
                   feature_limit         = 26,
                   character_limit       = 30,
                   show_heatmap_col_name = FALSE,
                   show_col              = FALSE,
                   show_plot             = TRUE)
#> [1] ">>>  Processing signature: tumor_signature"
```

```
#> [1] ">>>  Processing signature: EMT"
```

```
head(res)
#> # A tibble: 6 x 6
#>   sig_names                         p.value statistic    p.adj log10pvalue stars
#>   <chr>                               <dbl>     <dbl>    <dbl>       <dbl> <fct>
#> 1 EMT3                             1.34e-21    -0.482 2.14e-20       20.9  **** 
#> 2 Positive_regulation_of_exosomal… 9.69e-18     0.438 7.75e-17       17.0  **** 
#> 3 EMT2                             4.30e-17    -0.430 2.29e-16       16.4  **** 
#> 4 EV_Cell_2020                     6.95e-16    -0.415 2.78e-15       15.2  **** 
#> 5 Nature_metabolism_Hypoxia        7.70e-15     0.401 2.46e-14       14.1  **** 
#> 6 Ferroptosis                      2.27e- 6    -0.250 6.05e- 6        5.64 ****
```

###### 3) Evaluate Pan-F-TBRs related TME cell infiltration

Analyze the immune cell infiltration landscape related to Pan-F-TBRs.

```
names(sig_group)
#>  [1] "tumor_signature"        "EMT"                    "io_biomarkers"         
#>  [4] "immu_microenvironment"  "immu_suppression"       "immu_exclusion"        
#>  [7] "immu_exhaustion"        "TCR_BCR"                "tme_signatures1"       
#> [10] "tme_signatures2"        "Bcells"                 "Tcells"                
#> [13] "DCs"                    "Macrophages"            "Neutrophils"           
#> [16] "Monocytes"              "CAFs"                   "NK"                    
#> [19] "tme_cell_types"         "CIBERSORT"              "MCPcounter"            
#> [22] "EPIC"                   "xCell"                  "quanTIseq"             
#> [25] "ESTIMATE"               "IPS"                    "TIMER"                 
#> [28] "fatty_acid_metabolism"  "hypoxia_signature"      "cholesterol_metabolism"
#> [31] "Metabolism"             "hallmark"               "hallmark1"             
#> [34] "hallmark2"              "hallmark3"              "Rooney_et_al"          
#> [37] "Bindea_et_al"           "Li_et_al"               "Peng_et_al"
res<-iobr_cor_plot(pdata_group           = pdata_group,
                   id1                   = "ID",
                   feature_data          = imvigor210_sig,
                   id2                   = "ID",
                   target                = "Pan_F_TBRs",
                   group                 = "group3",
                   is_target_continuous  = TRUE,
                   padj_cutoff           = 1,
                   index                 = 6,
                   category              = "signature",
                   signature_group       = sig_group[20:24],
                   ProjectID             = "IMvigor210",
                   palette_box           = "jco",
                   palette_corplot       = "pheatmap",
                   palette_heatmap       = 3,
                   feature_limit         = 26,
                   character_limit       = 30,
                   show_heatmap_col_name = FALSE,
                   show_col              = FALSE,
                   show_plot             = TRUE)
#> [1] ">>>  Processing signature: CIBERSORT"
```

```
#> [1] ">>>  Processing signature: MCPcounter"
```

```
#> [1] ">>>  Processing signature: EPIC"
```

```
#> [1] ">>>  Processing signature: xCell"
```

```
#> [1] ">>>  Processing signature: quanTIseq"
```

```
head(res)
#> # A tibble: 6 x 6
#>   sig_names               p.value statistic    p.adj log10pvalue stars
#>   <chr>                     <dbl>     <dbl>    <dbl>       <dbl> <fct>
#> 1 Epithelial_cells_xCell 8.39e-42     0.642 9.91e-40        41.1 **** 
#> 2 Sebocytes_xCell        1.31e-26     0.530 7.71e-25        25.9 **** 
#> 3 Astrocytes_xCell       8.94e-26    -0.523 3.52e-24        25.0 **** 
#> 4 StromaScore_xCell      7.31e-24    -0.504 2.16e-22        23.1 **** 
#> 5 Fibroblasts_xCell      1.66e-23    -0.501 3.92e-22        22.8 **** 
#> 6 Keratinocytes_xCell    7.31e-21     0.474 1.44e-19        20.1 ****
```

##### 5.2.5 Batch Analyses and Visualization

IOBR provide multiple batch analytic functions for statistical analysis, data operation and transformation. For instance,batch survival analysis could be employed for batch analysis of the best cutoff value for subsequent batch survival analysis, varied batch statistical tests, etc.

###### 1) Batch survival analyses

```
input<-imvigor210_pdata %>% 
  filter(!is.na(.$OS_days)) %>% 
  filter(!is.na(.$OS_status)) %>% 
  mutate(OS_days = as.numeric(.$OS_days)) %>% 
  mutate(OS_status = as.numeric(.$OS_status)) %>% 
  dplyr::select(ID,OS_days,OS_status) %>% 
  inner_join(.,imvigor210_sig, by= "ID")

res <- batch_surv(pdata    = input, 
                  variable = c(4:ncol(input)), 
                  time     = "OS_days",
                  status   = "OS_status")
#> Warning in fitter(X, Y, istrat, offset, init, control, weights = weights, :
#> Loglik converged before variable 1 ; coefficient may be infinite.
head(res)
#> # A tibble: 6 x 5
#>   ID                                P       HR CI_low_0.95 CI_up_0.95
#>   <chr>                         <dbl>    <dbl>       <dbl>      <dbl>
#> 1 Hepatocytes_xCell          0        Inf           78.8     Inf     
#> 2 IFNG_signature_Ayers_et_al 0.0001     0.814        0.733     0.905 
#> 3 Macrophages_M1_CIBERSORT   0.0002     0.0014       0         0.0421
#> 4 Lysine_Degradation         0.0004     1.32         1.13      1.53  
#> 5 TMEscoreA_plus             0.000600   0.886        0.827     0.950 
#> 6 CD_8_T_effector            0.0008     0.856        0.781     0.937
```

###### 2) Subgroup survival analyses

```
##loading data and filter NA
data(subgroup_data)
input <- subgroup_data %>% 
   filter(time > 0) %>% 
   filter(!is.na(status)) %>% 
   filter(!is.na(AJCC_stage))

##for binary variable
res <- subgroup_survival(pdata    = input,
                         time     = "time", 
                         status   = "status",
                         variable = c("ProjectID", "AJCC_stage"),
                         object   = "score_binary" )
res
#>                         P     HR CI_low_0.95 CI_up_0.95
#> ProjectID_Dataset1 0.0000 3.0901      1.8648     5.1205
#> ProjectID_Dataset2 0.0023 2.1049      1.3057     3.3931
#> ProjectID_Dataset3 0.1397 2.4431      0.7467     7.9940
#> ProjectID_Dataset4 0.0024 2.0312      1.2854     3.2097
#> ProjectID_Dataset5 0.0000 4.6375      2.3759     9.0520
#> AJCC_stage_1       0.2533 1.7307      0.6753     4.4354
#> AJCC_stage_2       0.0000 2.9450      1.7858     4.8569
#> AJCC_stage_3       0.0000 2.2821      1.5551     3.3490
#> AJCC_stage_4       0.0510 1.5941      0.9979     2.5466
```

###### 3) Batch statistical analyses

```
data("imvigor210_sig")
data("imvigor210_pdata")
# For analyses of binary variables
input<-merge(imvigor210_pdata[,c("ID","BOR_binary")],imvigor210_sig,by="ID",all = F)
input<-input[!input$BOR_binary=="NA",]
input[1:5,1:5]
#>                ID BOR_binary B_cells_naive_CIBERSORT B_cells_memory_CIBERSORT
#> 2 SAM0257bbbbd388         NR              0.00000000               0.03161044
#> 3 SAM025b45c27e05         NR              0.00000000               0.00000000
#> 4 SAM032c642382a7         NR              0.08022624               0.00000000
#> 6 SAM0571f17f4045         NR              0.31954131               0.13731249
#> 7 SAM065890737112          R              0.10875636               0.00000000
#>   Plasma_cells_CIBERSORT
#> 2             0.03799257
#> 3             0.00167134
#> 4             0.01413286
#> 6             0.00000000
#> 7             0.01061609
res<-batch_wilcoxon(data    = input,
                    target  = "BOR_binary", 
                    group_names = c("NR","R"),
                    feature = colnames(imvigor210_sig)[3:ncol(imvigor210_sig)])
head(res, n=10)
#> # A tibble: 10 x 8
#>    sig_names         p.value      NR       R statistic   p.adj log10pvalue stars
#>    <chr>               <dbl>   <dbl>   <dbl>     <dbl>   <dbl>       <dbl> <fct>
#>  1 Mismatch_Repair   9.21e-6 -0.111   0.246    -0.357  0.00138        5.04 **** 
#>  2 Cell_cycle        1.05e-5 -0.297   0.673    -0.969  0.00138        4.98 **** 
#>  3 TMEscoreB_plus    1.35e-5  0.149  -0.432     0.581  0.00138        4.87 **** 
#>  4 DNA_replication   1.38e-5 -0.147   0.353    -0.500  0.00138        4.86 **** 
#>  5 DDR               1.52e-5 -0.229   0.532    -0.760  0.00138        4.82 **** 
#>  6 Nucleotide_excis… 2.16e-5 -0.106   0.244    -0.350  0.00154        4.67 **** 
#>  7 Homologous_recom… 3.31e-5 -0.118   0.272    -0.390  0.00154        4.48 **** 
#>  8 HALLMARK_E2F_TAR… 3.33e-5  0.231   0.256    -0.0251 0.00154        4.48 **** 
#>  9 T_cells_follicul… 3.48e-5  0.0164  0.0380   -0.0216 0.00154        4.46 **** 
#> 10 HALLMARK_WNT_BET… 3.70e-5  0.144   0.132     0.0119 0.00154        4.43 ****
#Note: Features with statistically significant p value could be used to construct model in the next module.
model_feas<-as.character(res[res$p.value<0.05,"sig_names"])

# For analyses of continuous variables
# head(imvigor210_sig)
res<-batch_cor(data    = imvigor210_sig,
               target  = "Pan_F_TBRs",
               feature = colnames(imvigor210_sig)[3:ncol(imvigor210_sig)],
               method  = "spearman")
head(res, n=10)
#> # A tibble: 10 x 6
#>    sig_names                       p.value statistic     p.adj log10pvalue stars
#>    <chr>                             <dbl>     <dbl>     <dbl>       <dbl> <fct>
#>  1 Normal_mucosa_Bindea_et_al    2.28e-146     0.924 1.03e-143        146. **** 
#>  2 TMEscoreB_CIR                 6.06e-145     0.922 1.37e-142        144. **** 
#>  3 Fibroblasts_MCPcounter        1.07e-144     0.922 1.62e-142        144. **** 
#>  4 CAF_Peng_et_al                2.48e-127     0.901 2.81e-125        127. **** 
#>  5 EMT2                          7.72e-124     0.896 7.00e-122        123. **** 
#>  6 HALLMARK_EPITHELIAL_MESENCHY… 1.63e-123     0.895 1.23e-121        123. **** 
#>  7 HALLMARK_MYOGENESIS           2.02e-113     0.879 1.31e-111        113. **** 
#>  8 HALLMARK_APICAL_JUNCTION      8.79e-112     0.876 4.98e-110        111. **** 
#>  9 TMEscoreB_plus                2.82e-106     0.866 1.42e-104        106. **** 
#> 10 StromalScore_estimate         1.21e-104     0.863 5.46e-103        104. ****
```

###### 4) Batch analyses of Pearson’s correlation coefficient

Detail code to perform `batch_pcc()`function for batch analyses of Partial Correlation coefficient(PCC) is shown below.

Herein, we utilized the data of the Pan\_F\_TBRs relevant signatures after adjusting TumorPurity as an example input. The major limitation of this method is that it may merely suitable for correlation estimation between two continuous variables.

```
pdata_group <- imvigor210_pdata[, c("ID", "TumorPurity", "Pan_F_TBRs")] %>% 
  rename(target = Pan_F_TBRs) %>% mutate(target = as.numeric(target))
res <- batch_pcc(pdata_group    = pdata_group, 
                 id1            = "ID", 
                 feature_data   = imvigor210_sig, 
                 id2            = "ID", 
                 interferenceid = "TumorPurity",
                 target         = "target",
                 method         = "pearson")
head(res, n = 8)
#> # A tibble: 8 x 6
#>   sig_names                         p.value statistic    p.adj log10pvalue stars
#>   <chr>                               <dbl>     <dbl>    <dbl>       <dbl> <fct>
#> 1 Epithelial_cells_xCell           1.26e-25     0.522 5.72e-23        24.9 **** 
#> 2 Keratinocytes_xCell              1.24e-16     0.425 2.82e-14        15.9 **** 
#> 3 MHC_Class_I                      6.63e-15     0.402 1.01e-12        14.2 **** 
#> 4 APM                              1.78e-13     0.382 2.03e-11        12.7 **** 
#> 5 Sebocytes_xCell                  2.38e-13     0.380 2.17e-11        12.6 **** 
#> 6 Riboflavin_Metabolism            9.02e-12     0.355 6.84e-10        11.0 **** 
#> 7 HALLMARK_PI3K_AKT_MTOR_SIGNALING 1.13e-11     0.354 7.35e-10        10.9 **** 
#> 8 HALLMARK_P53_PATHWAY             1.98e-11     0.350 1.13e- 9        10.7 ****
```

#### 5.3 Mutation Module

##### 5.3.1 Load Mutation MAF data

The input Mutation matrix with MAF data format is loaded for further analyses.

MAF data of TCGA-STAD was obtained from UCSC Xena website

```
# help("make_mut_matrix")

maf_file<-"TCGA.STAD.mutect.c06465a3-50e7-46f7-b2dd-7bd654ca206b.DR-10.0.somatic.maf"
mut_list<-make_mut_matrix(maf      = maf_file,
                          isTCGA   = T, 
                          category = "multi")
#> -Reading
#> -Validating
#> --Removed 2 duplicated variants
#> -Silent variants: 70966 
#> -Summarizing
#> --Possible FLAGS among top ten genes:
#>   TTN
#>   MUC16
#>   SYNE1
#>   FLG
#> -Processing clinical data
#> --Missing clinical data
#> -Finished in 13.8s elapsed (22.1s cpu) 
#>        Frame_Shift_Del        Frame_Shift_Ins           In_Frame_Del 
#>                  18418                   4461                    692 
#>           In_Frame_Ins      Missense_Mutation      Nonsense_Mutation 
#>                    268                 109668                   6011 
#>       Nonstop_Mutation            Splice_Site Translation_Start_Site 
#>                    107                   2445                    106 
#>    DEL    INS    SNP 
#>  19387   4900 117889
#> Aggregation function missing: defaulting to length
#> Aggregation function missing: defaulting to length
#> Aggregation function missing: defaulting to length
#> Aggregation function missing: defaulting to length
# If "multi" is set in above "category" parameter, four data frames will be returned, which evaluate all the mutations of every gene or estimate only SNP, indel and frameshift as follow:

# NOTE: UCSCXenaTools can be applied to access variant data of TCGA data sets.
var_stad<-XenaGenerate(subset = XenaCohorts =="GDC TCGA Stomach Cancer (STAD)") %>% 
  XenaFilter(filterDatasets    = "TCGA-STAD.mutect2_snv.tsv") %>% 
  XenaQuery() %>%
  XenaDownload() %>% 
  XenaPrepare()
#> This will check url status, please be patient.
#> All downloaded files will under directory /tmp/Rtmpmi7SbB.
#> The 'trans_slash' option is FALSE, keep same directory structure as Xena.
#> Creating directories for datasets...
#> Downloading TCGA-STAD.mutect2_snv.tsv.gz
head(var_stad)
#> # A tibble: 6 x 11
#>   Sample_ID gene  chrom  start    end ref   alt   Amino_Acid_Chan… effect filter
#>   <chr>     <chr> <chr>  <dbl>  <dbl> <chr> <chr> <chr>            <chr>  <chr> 
#> 1 TCGA-CD-… C1or… chr1  2.19e6 2.19e6 G     -     p.P72Rfs*87      frame… PASS  
#> 2 TCGA-CD-… ERRF… chr1  8.01e6 8.01e6 C     T     p.P327P          synon… panel…
#> 3 TCGA-CD-… CLCN6 chr1  1.18e7 1.18e7 G     A     p.S486N          misse… PASS  
#> 4 TCGA-CD-… PRAM… chr1  1.28e7 1.28e7 G     A     p.G341R          misse… panel…
#> 5 TCGA-CD-… PRAM… chr1  1.32e7 1.32e7 G     T     p.P148H          misse… PASS  
#> 6 TCGA-CD-… CELA… chr1  1.55e7 1.55e7 G     A     p.G59R           misse… PASS  
#> # … with 1 more variable: dna_vaf <dbl>
# Then function `make_mut_matrix` can be used to transform data frame into mutation matrix
mut_list2<-make_mut_matrix(mut_data               = var_stad,
                           category               = "multi",
                           Tumor_Sample_Barcode   = "Sample_ID",
                           Hugo_Symbol            = "gene",
                           Variant_Classification = "effect",
                           Variant_Type           = "Variant_Type")
#> Aggregation function missing: defaulting to length
#> Aggregation function missing: defaulting to length
#> Aggregation function missing: defaulting to length
#> Aggregation function missing: defaulting to length

# NOTE: IOBR provides mutation matrix(`tcga_stad_var`), if MAF data or UCSC can not be accessed. 
tcga_stad_var[1:5,1:5]
#>              ABCA12 ABCA13 ABCA2 ABCB1 ABCC9
#> TCGA-3M-AB46      0      0     0     0     0
#> TCGA-3M-AB47      0      0     0     0     0
#> TCGA-B7-5816      0      2     0     1     1
#> TCGA-B7-5818      0      0     0     0     0
#> TCGA-B7-A5TI      0      0     0     1     0
```

##### 5.3.2 Analyze Signature Associated Mutations

Utilize `sig_group()` function for batch analyses and visualization of response associated signatures.

```
names(mut_list)
#> [1] "all"        "snp"        "indel"      "frameshift"
mut_list$all[1:5, 1:10]
#>              A1BG A1CF A2M A2ML1 A3GALT2 A4GALT A4GNT AAAS AACS AADAC
#> TCGA-3M-AB46    1    1   0     0       0      0     0    0    0     0
#> TCGA-3M-AB47    0    0   0     0       0      0     0    0    0     0
#> TCGA-B7-5816    0    0   1     0       0      0     0    1    0     0
#> TCGA-B7-5818    0    0   0     0       0      0     0    0    0     0
#> TCGA-B7-A5TI    0    0   0     0       0      0     0    0    0     0

# choose SNP mutation matrix as input
mut<-mut_list$snp
# NOTE: If the maximum of mutation counts of a single gene of a person in is over 4, the mutation data would be standardized according to following principles.
# mut[mut>=3&mut<=5]<-3
# mut[mut>5]<-4
# Each gene of every samples is categorized as binary variables(mutation or non-mutation) in the MAF data. 
##########################
res<-find_mutations(mutation_matrix     = mut, 
                    signature_matrix    = tcga_stad_sig,
                    id_signature_matrix = "ID",
                    signature           = "CD_8_T_effector",
                    min_mut_freq        = 0.01,
                    plot                = TRUE,
                    method              = "Wilcoxon",
                    save_path           = paste0("CD_8_T_effector-relevant-mutations"),
                    palette             = "jco",
                    show_plot           = T)
#> [1] ">>>> Result of Wilcoxon test"
#>             p.value  names statistic adjust_pvalue
#> PIK3CA 1.921035e-10 PIK3CA      4125  9.605174e-08
#> TCHH   1.961642e-05   TCHH      3312  4.904106e-03
#> SPEG   3.532750e-05   SPEG      1947  5.887916e-03
#> LRP1   7.511741e-05   LRP1      2649  9.389676e-03
#> WDFY3  1.257659e-04  WDFY3      2964  1.257659e-02
#> ARID1A 2.468609e-04 ARID1A      4878  2.057174e-02
#> PLXNA4 4.215809e-04 PLXNA4      3638  3.011292e-02
#> ANK3   6.399572e-04   ANK3      4446  3.933979e-02
#> DMD    7.364591e-04    DMD      5311  3.933979e-02
#> PLEC   8.026240e-04   PLEC      5562  3.933979e-02
```

#### 5.4 Model Construction Module

For effective application of the signatures in clinical interpretation, IOBR also provides functions for feature selection, robust biomarker identification, and model construction based on priorly identified phenotype associated biomarkers in 5.2.1 Explore Phenotype Relevant Signatures. For predictive model construction.

```
data("imvigor210_sig")
data("imvigor210_pdata")
# For analyses of binary variables
input<-imvigor210_pdata %>% 
  dplyr::select(ID,BOR_binary) %>% 
  inner_join(.,imvigor210_sig,by="ID") %>% 
  filter(!is.na(.$BOR_binary)) %>% 
  filter(!.$BOR_binary=="NA")

# Feature engineering
res<-batch_wilcoxon(data    = as.data.frame(input),
                    target  = "BOR_binary",
                    group_names = c("NR","R"),
                    feature = colnames(input)[3:ncol(input)])
head(res)
#> # A tibble: 6 x 8
#>   sig_names            p.value     NR      R statistic   p.adj log10pvalue stars
#>   <chr>                  <dbl>  <dbl>  <dbl>     <dbl>   <dbl>       <dbl> <fct>
#> 1 Mismatch_Repair      9.21e-6 -0.111  0.246    -0.357 0.00138        5.04 **** 
#> 2 Cell_cycle           1.05e-5 -0.297  0.673    -0.969 0.00138        4.98 **** 
#> 3 TMEscoreB_plus       1.35e-5  0.149 -0.432     0.581 0.00138        4.87 **** 
#> 4 DNA_replication      1.38e-5 -0.147  0.353    -0.500 0.00138        4.86 **** 
#> 5 DDR                  1.52e-5 -0.229  0.532    -0.760 0.00138        4.82 **** 
#> 6 Nucleotide_excisio…  2.16e-5 -0.106  0.244    -0.350 0.00154        4.67 ****
model_feas<-as.character(res[res$p.value<0.05,]$sig_names)

input<-as.data.frame(imvigor210_sig)
feas<-colnames(input)[colnames(input)%in%model_feas]
input<-input[, c("ID",feas)]

# target data
pdata_group <- imvigor210_pdata[!imvigor210_pdata$BOR_binary=="NA",c("ID","BOR_binary")]
pdata_group$BOR_binary <- ifelse(pdata_group$BOR_binary == "R", 1, 0)

#Feature selection
binomial_result <- BinomialModel(x           = input, 
                                 y           = pdata_group, 
                                 seed        = "123456", 
                                 scale       = TRUE,
                                 train_ratio = 0.7, 
                                 nfold       = 10, 
                                 plot        = T)
#> NULL
#> NULL
#> NULL
#> NULL
```

```
#> NULL

plot(binomial_result$lasso_result$model)
```

```
plot(binomial_result$ridge_result$model)
```

```
lapply(binomial_result[1:2], function(z)z$AUC)
#> $lasso_result
#>       lambda.min lambda.1se
#> train  0.8911641        0.5
#> test   0.7017974        0.5
#> 
#> $ridge_result
#>       lambda.min lambda.1se
#> train  0.7915115        0.5
#> test   0.7263072        0.5
```

For prognostic model construction.

```
data("imvigor210_pdata")
pdata_prog<-imvigor210_pdata %>% 
  dplyr::select(ID, OS_days, OS_status) %>% 
  filter(!is.na(.$OS_days)) %>% 
  filter(!.$OS_days=="NA") %>% 
  dplyr:: rename(time = OS_days) %>% 
  dplyr:: rename(status = OS_status) %>% 
  mutate(time = as.numeric(.$time)) %>%
  mutate(status = as.numeric(.$status))

pdata_prog<-as.data.frame(pdata_prog)
input<-as.data.frame(imvigor210_sig)

prognostic_result <- PrognosticModel(x           = input, 
                                     y           = pdata_prog, 
                                     scale       = T, 
                                     seed        = "123456", 
                                     train_ratio = 0.8,
                                     nfold       = 10,
                                     plot        = T)
#> NULL
#> NULL
#> NULL
```

```
plot(prognostic_result$lasso_result$model)
```

```
plot(prognostic_result$ridge_result$model)
```

In order to find best predictive or prognostic signature, we recommend Blasso package to determine the most robust markers. If Blasso R package is utilized in your published research, please cite: DQ Zeng, ZL Ye, et al., Macrophage correlates with immunophenotype and predicts anti-PD-L1 response of urothelial cancer. Theranostics 2020; 10(15):7002-7014. doi:10.7150/thno.46176

```
if (!requireNamespace("Blasso", quietly = TRUE))  
  devtools::install_github("DongqiangZeng0808/Blasso")
library("Blasso")
#For prognostic biomarkers
# res<-best_predictor_cox(target_data        = target, 
#                         features           = features, 
#                         status             = "status",
#                         time               = "time",
#                         nfolds             = 10,
#                         permutation        = 300,
#                         show_progress      = FALSE)
#For predictive biomarkers
# res<-best_predictor_binomial(target_data   = target, 
#                              features      = features,
#                              response      = "status",
#                              nfolds        = 10,
#                              permutation   = 300,
#                              show_progress = FALSE)
```

### 6. Summary

IOBR provides four major analytic modules allowing effective and flexible analysis of tumor immunologic, clinical, genomics, and single-cell data. There might be some limitations in this package, we will put our efforts into its improvement for more functionality in the future. Any questions, bug reports, or suggestions about IOBR R package is appreciated. Please contact us through E-mail to.

### 7. Session Information

```
sessionInfo()
#> R version 3.6.3 (2020-02-29)
#> Platform: x86_64-pc-linux-gnu (64-bit)
#> Running under: CentOS Linux 7 (Core)
#> 
#> Matrix products: default
#> BLAS/LAPACK: /usr/lib64/libopenblasp-r0.3.3.so
#> 
#> locale:
#>  [1] LC_CTYPE=en_US.UTF-8 LC_NUMERIC=C         LC_TIME=C           
#>  [4] LC_COLLATE=C         LC_MONETARY=C        LC_MESSAGES=C       
#>  [7] LC_PAPER=C           LC_NAME=C            LC_ADDRESS=C        
#> [10] LC_TELEPHONE=C       LC_MEASUREMENT=C     LC_IDENTIFICATION=C 
#> 
#> attached base packages:
#> [1] parallel  stats     graphics  grDevices utils     datasets  methods  
#> [8] base     
#> 
#> other attached packages:
#>  [1] Blasso_0.1.0        progress_1.2.2      RColorBrewer_1.1-2 
#>  [4] glmnet_4.0-2        Matrix_1.2-18       GEOquery_2.54.1    
#>  [7] Biobase_2.46.0      BiocGenerics_0.32.0 UCSCXenaTools_1.3.4
#> [10] maftools_2.2.10     tidyHeatmap_1.1.5   forcats_0.5.0      
#> [13] stringr_1.4.0       purrr_0.3.4         readr_1.3.1        
#> [16] tidyr_1.1.0         tidyverse_1.3.0     MCPcounter_1.2.0   
#> [19] curl_4.3            estimate_1.0.13     EPIC_1.1.5         
#> [22] IOBR_0.99.9         survival_3.2-7      ggpubr_0.4.0       
#> [25] ggplot2_3.3.2       dplyr_1.0.0         tibble_3.0.1       
#> 
#> loaded via a namespace (and not attached):
#>   [1] utf8_1.1.4                  tidyselect_1.1.0           
#>   [3] RSQLite_2.2.0               AnnotationDbi_1.48.0       
#>   [5] htmlwidgets_1.5.1           grid_3.6.3                 
#>   [7] BiocParallel_1.20.1         lpSolve_5.6.15             
#>   [9] munsell_0.5.0               pec_2019.11.03             
#>  [11] codetools_0.2-16            preprocessCore_1.48.0      
#>  [13] withr_2.2.0                 colorspace_1.4-1           
#>  [15] limSolve_1.5.6              knitr_1.29                 
#>  [17] rstudioapi_0.11             ROCR_1.0-11                
#>  [19] stats4_3.6.3                ggsignif_0.6.0             
#>  [21] labeling_0.3                GenomeInfoDbData_1.2.2     
#>  [23] KMsurv_0.1-5                farver_2.0.3               
#>  [25] bit64_0.9-7                 vctrs_0.3.1                
#>  [27] generics_0.1.0              xfun_0.15                  
#>  [29] BiocFileCache_1.10.2        R6_2.4.1                   
#>  [31] GenomeInfoDb_1.22.1         clue_0.3-57                
#>  [33] timereg_1.9.8               locfit_1.5-9.4             
#>  [35] bitops_1.0-6                DelayedArray_0.12.3        
#>  [37] assertthat_0.2.1            promises_1.1.1             
#>  [39] scales_1.1.1                nnet_7.3-12                
#>  [41] gtable_0.3.0                sva_3.34.0                 
#>  [43] rlang_0.4.6                 genefilter_1.68.0          
#>  [45] GlobalOptions_0.1.2         splines_3.6.3              
#>  [47] rstatix_0.6.0               acepack_1.4.1              
#>  [49] wordcloud_2.6               broom_0.5.6                
#>  [51] checkmate_2.0.0             reshape2_1.4.4             
#>  [53] yaml_2.2.1                  abind_1.4-5                
#>  [55] modelr_0.1.8                backports_1.1.8            
#>  [57] httpuv_1.5.4                Hmisc_4.4-0                
#>  [59] lava_1.6.8.1                tools_3.6.3                
#>  [61] ellipsis_0.3.1              plyr_1.8.6                 
#>  [63] Rcpp_1.0.5                  base64enc_0.1-3            
#>  [65] zlibbioc_1.32.0             RCurl_1.98-1.2             
#>  [67] prettyunits_1.1.1           rpart_4.1-15               
#>  [69] openssl_1.4.2               GetoptLong_1.0.4           
#>  [71] viridis_0.5.1               cowplot_1.0.0              
#>  [73] S4Vectors_0.24.4            zoo_1.8-8                  
#>  [75] SummarizedExperiment_1.16.1 haven_2.3.1                
#>  [77] cluster_2.1.0               fs_1.4.1                   
#>  [79] magrittr_1.5                data.table_1.12.8          
#>  [81] openxlsx_4.1.5              circlize_0.4.11            
#>  [83] reprex_0.3.0                survminer_0.4.8            
#>  [85] mvtnorm_1.1-1               timeROC_0.4                
#>  [87] matrixStats_0.56.0          hms_0.5.3                  
#>  [89] mime_0.9                    evaluate_0.14              
#>  [91] GSVA_1.34.0                 xtable_1.8-4               
#>  [93] XML_3.99-0.3                rio_0.5.16                 
#>  [95] jpeg_0.1-8.1                readxl_1.3.1               
#>  [97] IRanges_2.20.2              gridExtra_2.3              
#>  [99] shape_1.4.5                 compiler_3.6.3             
#> [101] biomaRt_2.42.1              crayon_1.3.4               
#> [103] htmltools_0.5.0             mgcv_1.8-31                
#> [105] later_1.1.0.1               Formula_1.2-3              
#> [107] geneplotter_1.64.0          lubridate_1.7.9            
#> [109] DBI_1.1.0                   corrplot_0.84              
#> [111] ppcor_1.1                   dbplyr_1.4.4               
#> [113] ComplexHeatmap_2.2.0        rappdirs_0.3.1             
#> [115] MASS_7.3-51.5               car_3.0-8                  
#> [117] cli_2.0.2                   quadprog_1.5-8             
#> [119] GenomicRanges_1.38.0        pkgconfig_2.0.3            
#> [121] km.ci_0.5-2                 prettydoc_0.4.0            
#> [123] numDeriv_2016.8-1.1         foreign_0.8-75             
#> [125] xml2_1.3.2                  foreach_1.5.0              
#> [127] annotate_1.64.0             ggcorrplot_0.1.3           
#> [129] XVector_0.26.0              prodlim_2019.11.13         
#> [131] rvest_0.3.5                 digest_0.6.25              
#> [133] pracma_2.2.9                graph_1.64.0               
#> [135] rmarkdown_2.3               cellranger_1.1.0           
#> [137] survMisc_0.5.5              htmlTable_2.0.0            
#> [139] GSEABase_1.48.0             shiny_1.5.0                
#> [141] rjson_0.2.20                lifecycle_0.2.0            
#> [143] nlme_3.1-144                jsonlite_1.7.0             
#> [145] carData_3.0-4               viridisLite_0.3.0          
#> [147] askpass_1.1                 limma_3.42.2               
#> [149] fansi_0.4.1                 pillar_1.4.4               
#> [151] ggsci_2.9                   lattice_0.20-38            
#> [153] fastmap_1.0.1               httr_1.4.1                 
#> [155] glue_1.4.1                  remotes_2.1.1              
#> [157] zip_2.0.4                   png_0.1-7                  
#> [159] shinythemes_1.1.2           iterators_1.0.12           
#> [161] bit_1.1-15.2                class_7.3-15               
#> [163] stringi_1.4.6               blob_1.2.1                 
#> [165] DESeq2_1.26.0               latticeExtra_0.6-29        
#> [167] memoise_1.1.0               e1071_1.7-4
```

### 8. Citing IOBR

In that IOBR incorporating some algorithms and methodologies from many other packages, please cite corresponding papers alongside IOBR when using the functions contains these algorithms. The detailed paper sources and respective licenses were listed in the subset of 5.1.3 Available Methods to Decode TME Contexture.

If IOBR R package is utilized in your published research, please cite:

> - DQ Zeng, ZL Ye, RF Shen, Y Xiong, …, WJ Liao\*. (2020). IOBR: Multi-omics Immuno-Oncology Biological Research to decode tumor microenvironment and signatures. [doi……..]

Please cite the following papers appropriately for TME deconvolution algorithm if used:

> - CIBERSORT: Newman, A. M., Liu, C. L., Green, M. R., Gentles, A. J., Feng, W., Xu, Y., … Alizadeh, A. A. (2015). Robust enumeration of cell subsets from tissue expression profiles. Nature Methods, 12(5), 453–457. https://doi.org/10.1038/nmeth.3337
> - ESTIMATE: Vegesna R, Kim H, Torres-Garcia W, …, Verhaak R.\*(2013). Inferring tumour purity and stromal and immune cell admixture from expression data. Nature Communications 4, 2612. http://doi.org/10.1038/ncomms3612
> - quanTIseq: Finotello, F., Mayer, C., Plattner, C., Laschober, G., Rieder, D., Hackl, H., …, Sopper, S.\* (2019). Molecular and pharmacological modulators of the tumor immune contexture revealed by deconvolution of RNA-seq data. Genome medicine, 11(1), 34. https://doi.org/10.1186/s13073-019-0638-6
> - Li, B., Severson, E., Pignon, J.-C., Zhao, H., Li, T., Novak, J., … Liu, X. S.\* (2016). Comprehensive analyses of tumor immunity: implications for cancer immunotherapy. Genome Biology, 17(1), 174.
> - IPS: P. Charoentong et al.\*, Pan-cancer Immunogenomic Analyses Reveal Genotype-Immunophenotype Relationships and Predictors of Response to Checkpoint Blockade. Cell Reports 18, 248-262 (2017). https://doi.org/10.1016/j.celrep.2016.12.019
> - MCPCounter: Becht, E., Giraldo, N. A., Lacroix, L., Buttard, B., Elarouci, N., Petitprez, F., … de Reyniès, A\*. (2016). Estimating the population abundance of tissue-infiltrating immune and stromal cell populations using gene expression. Genome Biology, 17(1), 218. https://doi.org/10.1186/s13059-016-1070-5
> - xCell: Aran, D., Hu, Z., & Butte, A. J.\* (2017). xCell: digitally portraying the tissue cellular heterogeneity landscape. Genome Biology, 18(1), 220. https://doi.org/10.1186/s13059-017-1349-1
> - EPIC: Racle, J., de Jonge, K., Baumgaertner, P., Speiser, D. E., & Gfeller, D\*. (2017). Simultaneous enumeration of cancer and immune cell types from bulk tumor gene expression data. ELife, 6, e26476. https://doi.org/10.7554/eLife.26476

For signature score estimation, please cite corresponding literature below:

> - ssgsea: Barbie, D.A. et al (2009). Systematic RNA interference reveals that oncogenic KRAS-driven cancers require TBK1. Nature, 462(5):108-112.
> - gsva: Hänzelmann, S., Castelo, R. and Guinney, J. (2013). GSVA: Gene set variation analysis for microarray and RNA-Seq data. BMC Bioinformatics, 14(1):7.
> - zscore: Lee, E. et al (2008). Inferring pathway activity toward precise disease classification. PLoS Comp Biol, 4(11):e1000217.

For the datasets enrolled in IOBR, please cite the data sources:

> - UCSCXena: Wang et al.,et al (2019). The UCSCXenaTools R package: a toolkit for accessing genomics data from UCSC Xena platform, from cancer multi-omics to single-cell RNA-seq. Journal of Open Source Software, 4(40), 1627
> - TLSscore: Helmink BA, Reddy SM, Gao J, et al. B cells and tertiary lymphoid structures promote immunotherapy response. Nature. 2020 Jan;577(7791):549-555.
> - IMvigor210 immuntherapy cohort: Mariathasan S, Turley SJ, Nickles D, et al. TGFβ attenuates tumour response to PD-L1 blockade by contributing to exclusion of T cells. Nature. 2018 Feb 22;554(7693):544-548.
> - HCP5: Kulski, J.K. Long Noncoding RNA HCP5, a Hybrid HLA Class I Endogenous Retroviral Gene: Structure, Expression, and Disease Associations. Cells 2019, 8, 480.
> - HCP5: Li, Y., Jiang, T., Zhou, W. et al. Pan-cancer characterization of immune-related lncRNAs identifies potential oncogenic biomarkers. Nat Commun 11, 1000 (2020).
> - HCP5: Sun J, Zhang Z, Bao S, et alIdentification of tumor immune infiltration-associated lncRNAs for improving prognosis and immunotherapy response of patients with non-small cell lung cancerJournal for ImmunoTherapy of Cancer 2020;8:e000110.
> - LINC00657: Feng Q, Zhang H, Yao D, Chen WD, Wang YD. Emerging Role of Non-Coding RNAs in Esophageal Squamous Cell Carcinoma. Int J Mol Sci. 2019 Dec 30;21(1):258. doi: 10.3390/ijms21010258.
> - LINC00657: Qin X, Zhou M, Lv H, Mao X, Li X, Guo H, Li L, Xing H. Long noncoding RNA LINC00657 inhibits cervical cancer development by sponging miR-20a-5p and targeting RUNX3. Cancer Lett. 2020 Oct 28:S0304-3835(20)30578-4. doi: 10.1016/j.canlet.2020.10.044.
> - LINC00657: Zhang XM, Wang J, Liu ZL, Liu H, Cheng YF, Wang T. LINC00657/miR-26a-5p/CKS2 ceRNA network promotes the growth of esophageal cancer cells via the MDM2/p53/Bcl2/Bax pathway. Biosci Rep. 2020;40(6):BSR20200525.
> - TCGA.STAD: Cancer Genome Atlas Research Network. Comprehensive molecular characterization of gastric adenocarcinoma. Nature. 2014 Sep 11;513(7517):202-9. doi: 10.1038/nature13480.
> - TCGA.STAD MAF data: https://api.gdc.cancer.gov/data/c06465a3-50e7-46f7-b2dd-7bd654ca206b
